# The Choreography of Coupled Brains: Coordinated Movement and Multi-Scale Neural Synchrony in Musical Theater for Individuals with Developmental Disabilities

**DOI:** 10.64898/2026.08.31.748378

**Authors:** Zachary Moore, Noor Tasnim, Ryan Frank, Carolyn Jones, Vidhi Tripathi, Grace Nobriga, Sindhu Dhaveji, Mary Gahagan, Jessica Purevtugs, Jesse Newpol, Julia C. Basso

## Abstract

Performing together, which demands moving, singing, and coordinating in real time, is among the most social things people do, yet whether such coordination synchronizes not only behavior but brain activity, within and between performers, has rarely been examined during genuine performance. In this pilot study, we used mobile electroencephalography, including dual-brain hyperscanning of co-performing partners, to resolve neural change over a 16-week musical-theater program in adolescents and adults with developmental disabilities, the majority autistic. We examined four levels: behavior, spectral power, within-brain (source-space) connectivity, and between-brain synchrony, framing all estimates as effect sizes. Anxiety decreased from pre- to post-intervention, consistently across participant and parent reports. During performance, high-beta and gamma power and within-brain amplitude connectivity rose broadly across the cortex, with maximal effects seen in parieto-occipital regions. Between-brain coupling was similarly carried by amplitude co-fluctuation rather than phase; however, it was graded by the degree of interpersonal movement coordination (strongest when partners mirrored one another) and scaled with movement vigor. Local and within-brain changes were thus broad and state-like, whereas coupling between brains was specific to interpersonal coordination. Coordinated movement may organize neural synchrony across scales, positioning musical theater as a tractable model for studying the social brain in motion.

## INTRODUCTION

Successful social interaction requires the continuous, reciprocal alignment of two people’s attention, actions, and their underlying brain activity. Neuroscientific hyperscanning studies, which record two or more brains simultaneously, show that the degree of neural synchrony between interacting partners tracks cooperation, communication, and social connectedness(Czeszumski *et al*., 2020; Czeszumski *et al*., 2022). Additionally, within a single brain, synchronized oscillatory activity supports the large-scale neural integration that underlies flexible cognition and social behavior(Varela *et al*., 2001; Hari and Kujala, 2009; Fries, 2015). A unifying proposal, the Synchronicity Hypothesis of Dance(Basso *et al*., 2020), holds that rhythmic, coordinated, socially embedded movement enhances neural synchrony at two levels at once — within an individual’s brain (intra-brain) and between the brains of interacting partners (inter-brain) — casting movement as an organizer of multiscale neural synchrony across sensory, motor, rhythmic, cognitive, social, emotional, and creative domains.

This framework is especially relevant to autism spectrum disorder (ASD) and related developmental disabilities, which are characterized by differences in social responsiveness and in interpreting others’ emotions and intentions(Lord *et al*., 2018; Lord *et al*., 2020), and which is marked by altered neural coordination at multiple scales(Uddin *et al*., 2013; O’Reilly *et al*., 2017). Autistic individuals show reduced intra-brain synchrony, including decreased long-range temporal correlations in the high-gamma band(Jia *et al*., 2021) and diminished alpha coherence and suppression(Mathewson *et al*., 2012). Emerging hyperscanning evidence suggests that inter-brain synchrony may likewise be reduced during socially demanding interaction(Kruppa *et al*., 2020; Azhari *et al*., 2025; Li *et al*., 2025). Co-occurring anxiety is common and is closely tied to social withdrawal(Lai *et al*., 2019; Kerns *et al*., 2020), compounding these challenges. Therefore, we hypothesize that synchrony-promoting activity may help repattern such atypical dynamics, making movement-rich, socially interactive experience a compelling, and largely untested avenue for ASD and related developmental disabilities.

Performing-arts interventions (including music, movement, and theater) engage social, emotional, and sensorimotor systems simultaneously, and a growing literature links them to behavioral and, in some cases, neural change in ASD. Improvisational music therapy improves social communication and increases resting-state connectivity between auditory, subcortical, and fronto-motor networks in autistic children(Geretsegger *et al*., 2014; Sharda *et al*., 2018). Additionally, the peer-mediated SENSE Theatre® program demonstrates gains in social competence alongside event-related-potential evidence of enhanced memory for faces, an index of social salience(Corbett *et al*., 2016; Corbett *et al*., 2023; Corbett *et al*., 2025). Further, dance/movement approaches have been used to build bodily awareness and self–other differentiation(Mastrominico *et al*., 2018). Yet the neurophysiology of active, group-based performance itself remains poorly characterized, notably because conventional neuroimaging requires participants to be still and isolated, precisely the opposite of performance. Advances in mobile electroencephalography (EEG) now make it feasible to record cortical activity during naturalistic movement and social interaction(Stangl *et al*., 2023; Basso *et al*., 2026).

The present pilot study used mobile EEG, including dual-EEG hyperscanning of co-performing partners, to characterize neural engagement across a 16-week musical-theatre program that culminated in a live production. Musical theater is a natural extension of the dance-based synchronicity framework: it layers movement, speech, music, and coordinated social interaction into a single, richly interpersonal activity. We examined behavioral change together with three complementary layers of the EEG signal (spectral power, intra-brain connectivity, and inter-brain synchrony). For the connectivity layers, we distinguished phase-based coupling from amplitude-envelope coupling. Intra-brain measures are the more tractable target in a small sample; inter-brain synchrony, though more challenging to explore and estimate in a small sample, is the novel frontier this design uniquely probes and the level tied most directly to the interpersonal claims of the hypothesis.

We tested two behaviorally grounded predictions and pursued a set of planned neural aims. Behaviorally, we predicted that participation would reduce anxiety and enhance mood and social connection. At the neural level, we predicted that interactive performance and movement would be associated with changes in oscillatory power and intra-brain synchrony, particularly in the beta and gamma bands linked to higher-order cognitive and sensorimotor processing(Jensen *et al*., 2007; Engel and Fries, 2010). For inter-brain synchrony, we predicted that shared performance would elevate coupling in the alpha band, given prior links between alpha and social processing(Hu *et al*., 2018; Park *et al*., 2024). Given the pilot sample, all neural analyses were framed around effect-size estimation and hypothesis generation rather than confirmatory inference.

## METHODS

### Participants and study design

Participants were recruited as a convenience sample through STEP VA (www.stepva.org), a nonprofit providing performing-arts opportunities for individuals with disabilities; consistent with the program’s all-ages structure, they spanned 9–30 years. Participants took part in a 16-week musical-theatre program, which included 16 weekly one-hour rehearsals culminating in a live production of *You’re a Good Man, Charlie Brown* (STEP VA, Fredericksburg, VA). Before and after the program, participants completed self-report psychosocial questionnaires and parents/guardians completed parallel proxy-report versions. Following the final performance, a subset of the participants underwent mobile EEG in a single session using a hyperscanning configuration (simultaneous recording in small groups), enabling both intra- and inter-brain analyses. This protocol was approved by the Virginia Tech Institutional Review Board (protocol #23-015). All participants provided written informed assent and parent/guardian consent, and all procedures were conducted in accordance with the Declaration of Helsinki.

### Measures

Participants completed a battery targeting emotional, psychological, and social health: the Revised Children’s Anxiety and Depression Scale (RCADS)(Chorpita *et al*., 2000), the Piers-Harris Self-Concept Scale, Third Edition(Piers *et al*., 2018), the KIDSCREEN-52(Ravens-Sieberer *et al*., 2008), the Social Connectedness Scale–Revised (SCS-R)(Lee *et al*., 2001), and the Emotional Regulation Strategies for Artistic Creative Activities Scale (ERS-ACA)(Fancourt *et al*., 2019), together with open-ended qualitative items; full item content and psychometric properties are given in Supplementary Methods. RCADS Total Anxiety was designated *a priori* as the primary behavioral outcome. Because no single instrument was normed across the full 9– 30 age range, all analyses used raw summed scores and within-subject pre-to-post change rather than age-normed scores, so that each participant served as their own baseline; an adult RCADS validation(McKenzie *et al*., 2019) supports interpretation of the primary outcome across the age range.

### EEG acquisition and preprocessing

Mobile EEG was recorded with 32-channel wet-electrode caps (LiveAmp 32; Brain Products, Gilching, Germany) as participants completed, in a single session, a three-minute eyes-open resting baseline, guided-breathing and movement-mirroring preparation exercises, and three musical numbers from the production (“You’re a Good Man, Charlie Brown”, “Beethoven Day,” and “Happiness”). Preprocessing used EEGLAB(Delorme and Makeig, 2004): data were bandpass-filtered (1–55 Hz), cleaned of noisy channels and high-variance segments by Artifact Subspace Reconstruction(Chang *et al*., 2018), interpolated to restore removed channels, re-referenced to the common average, and decomposed by Adaptive Mixture Independent Component Analysis(Hsu *et al*., 2018), with components classified by ICLabel(Pion-Tonachini *et al*., 2019). Components were retained for power analyses if labeled >50% brain(Gonsisko *et al*., 2023) and for connectivity analyses, after removing those labeled >90% muscle, eye, heart, line-noise, or other artifact. Recordings were screened for rank sufficiency before source and connectivity estimation(Westner *et al*., 2022; Pellegrini *et al*., 2023). Full parameters including filter design, channel-rejection, Artifact Subspace Reconstruction (ASR) settings, Adaptive Mixture Independent Component Analysis (AMICA) configuration, interpolation, component thresholds, and rank screening, are given in Supplementary Methods. Together, ASR followed by ICA-based decomposition and automated component classification reflects current recommended practice for isolating neural activity from movement and other non-brain artifacts in mobile EEG(Gorjan *et al*., 2022; Klug *et al*., 2024).

### Statistical analysis

#### Overview

As a pilot, inference is framed around effect-size estimation: effect sizes and the directional, spatial, and cross-rater consistency of effects are treated as the primary evidence, with p-values reported for transparency. Neural inference used permutation testing throughout (10,000 permutations), with family-wise error (FWER) control by max-statistic permutation(Nichols and Holmes, 2002) where a pre-specified cell family was defined and Benjamini-Hochberg false-discovery-rate (BH-FDR)(Benjamini and Hochberg, 1995) control within test families; analyses outside a pre-specified family are labeled exploratory. Behavioral and neural outcomes are reported in parallel; given the pilot sample, we did not test brain– behavior associations (e.g., correlations between neural and symptom change), which would be severely underpowered and yield unstable, inflated estimates at this *n*.

#### Behavioral and qualitative

Pre-to-post change was tested with Wilcoxon signed-rank tests separately by rater, with the matched-pairs rank-biserial correlation as effect size and bootstrap 95% confidence intervals (10,000 resamples); BH-FDR was applied within each instrument’s family of subscales and totals. Open-ended responses were analyzed by reflexive thematic analysis(Braun and Clarke, 2006), coding inductively and comparing pre and post at the group level with matched-pair comparison as a secondary layer (Supplementary Methods).

#### Spectral power

For each condition × ROI × band cell, within-subject change from baseline was tested with a paired t-test (effect size *d*z). FWER was controlled across a 27-cell pre-specified fast-band family (temporal, parietal, and prefrontal ROIs × low-beta, high-beta, gamma × performance, mirroring, and breathing vs. baseline) by max-statistic sign-flip permutation; alpha-suppression analyses were exploratory.

#### Intra-brain connectivity

The primary test was a pre-specified high-frequency omnibus: each participant’s mean connectivity change across all 28 network pairs in high-beta and gamma, tested against zero, collapsing the network grid to one value per participant. Band and condition specificity, within-versus between-network decomposition, and leave-one-subject-out robustness were examined; individual network-pair effects were exploratory.

#### Inter-brain synchrony

For each measure, band, and condition, real dyads were compared with same-condition pseudo-dyads by permutation (Cohen’s *d*), and within-dyad performance-versus-baseline contrasts by paired permutation (*d*z). BH-FDR was applied across the primary family (three musical numbers × two focal bands × three zero-lag measures: coherence, phase-locking value, and amplitude-envelope correlation); two zero-lag-insensitive measures (imaginary coherence and orthogonalized cross-brain AEC) served as leakage-robust sensitivity analyses. Because pseudo-dyads reuse individuals and are therefore not mutually independent, permutation p-values are treated as approximate, with inference resting on effect sizes, leave-one-dyad-out stability, and the spatial consistency of effects across the montage.

For both intra- and inter-brain analyses, phase-based measures (PLV, imaginary coherence) index the consistency of cycle-by-cycle timing between signals, whereas amplitude-based measures (envelope correlation, orthogonalized AEC) index co-fluctuation in oscillatory power, the degree to which the strength of the rhythm rises and falls together, independent of phase alignment; we report both because they reflect partly distinct coupling mechanisms.

#### Sensitivity analyses

All primary neural effects were re-examined with age as a covariate and tested for dependence on accelerometric movement vigor, with power spectra additionally decomposed into aperiodic and periodic components (specparam)(Donoghue *et al*., 2020). Full procedures are given in Supplementary Methods.

### Software and data availability

Analyses used MATLAB 2025b and Python 3.9; source reconstruction and connectivity used EEGLAB with the DIPFIT, ICLabel, and ROIconnect plugins, inter-brain synchrony used HyPyP(Ayrolles *et al*., 2021) on MNE-Python, and spectral parameterization used specparam(Donoghue *et al*., 2020). Analysis code and the de-identified derived data required to reproduce all figures and statistics are openly available on GitHub (https://github.com/embodiedbrainlab/stepva). Owing to the sensitive nature of the data and the small, vulnerable sample, raw EEG recordings and individual-level behavioral data are not publicly available but can be obtained from the corresponding author on reasonable request under a data-use agreement and IRB approval.

## RESULTS

### Participants

The study enrolled 22 participants and 22 parents/guardians; nine dyads (18 participants) withdrew, leaving 26 completers. Twelve participants completed EEG testing (nine autistic, two with intellectual disability, one with a chromosomal condition), and seven completed both pre- and post-intervention behavioral surveys (six autistic, one with intellectual disability); six completed both the surveys and EEG (five autistic, one with intellectual disability). The sample was predominantly male (64%), largely White (64%) and non-Hispanic (77%), and spanned childhood through adulthood (9–30 years). A majority reported an ASD diagnosis (55%); WASI composite scores fell below the normative mean (FSIQ-4 M = 77.5; Table 1).

**Table 1.** Demographic, diagnostic, and cognitive characteristics of participants and parent/guardian respondents.

| Characteristic | Participants (n = 22) | Parents/Guardians (n = 22) |
| --- | --- | --- |
| **Study completion, n (% of 22)** |  |  |
| Pre-test survey | 13 (59) | 12 (55) |
| Post-test survey | 13 (59) | 13 (59) |
| Both surveys | 7 (32) | 7 (32) |
| EEG testing | 12 (55) | — |
| EEG and surveys | 6 (27) | — |
| **Age (years), range / M (SEM)** |  |  |
|  | 9–30 / 17.4 (1.22) | 33–62 / 46.17 (1.54) |
| **Sex assigned at birth, n (%)** |  |  |
| Female | 8 (36) | 18 (82) |
| Male | 14 (64) | 0 (0) |
| Undisclosed | 0 (0) | 4 (18) |
| **Race, n (%)** |  |  |
| Black | 1 (5) | 0 (0) |
| White | 14 (64) | 17 (77) |
| Other | 3 (14) | 1 (5) |
| Undisclosed | 4 (18) | 4 (18) |
| **Ethnicity, n (%)** |  |  |
| Hispanic | 1 (5) | 1 (5) |
| Non-Hispanic | 17 (77) | 17 (77) |
| Undisclosed | 4 (18) | 4 (18) |
| **Education, n (%)** |  |  |
| In elementary school | 1 (5) | — |
| In middle school | 6 (27) | — |
| In high school | 3 (14) | — |
| Finished high school | 6 (27) | — |
| Some college | 2 (9) | 5 (23) |
| Bachelor's degree | — | 7 (32) |
| Advanced degree | — | 6 (27) |
| Undisclosed | 4 (18) | 4 (18) |
| **Diagnosis, n (%)** |  |  |
| ASD | 12 (55) | — |
| Intellectual disability | 3 (14) | — |
| Chromosomal abnormality | 1 (5) | — |
| Undisclosed | 6 (27) | — |
| **Household annual income, n (%)** |  |  |
| \$50,000–\$99,999 | — | 4 (18) |
| \$100,000–\$149,999 | — | 6 (27) |
| \$150,000–\$199,999 | — | 5 (23) |
| \$200,000 and over | — | 3 (14) |
| Undisclosed | — | 4 (18) |
| **WASI cognitive subtests (T-scores), range / M (SEM)** | **n = 13** |  |
| Block Design | 23–58 / 36.2 (2.95) | — |
| Vocabulary | 20–52 / 36.5 (3.07) | — |
| Matrix Reasoning | 18 – 60 / 42.8 (2.62) | — |
| Similarities | 20 – 60 / 28.9 (3.55) | — |
| **WASI composites, range / M (SEM)** | **n = 13** |  |
| Verbal Comprehension | 45 – 110 / 79.5 (5.6) | — |
| Perceptual Reasoning | 56 – 112 / 82.6 (4.75) | — |
| Full Scale IQ | 48 – 102 / 80.8 (4.4) | — |

**Note.** Values are n (%) of total enrolled (n = 22 in each column) unless otherwise noted. Em dash (—) indicates not applicable. M = mean; SEM = standard error of the mean. Reported diagnoses are participant/guardian reports confirmed through medical paperwork; six participants did not disclose a diagnosis. Twelve participants completed EEG testing; seven completed both pre- and post-intervention behavioral surveys; six completed both surveys and EEG testing. WASI-II = Wechsler Abbreviated Scale of Intelligence, Second Edition; subtests administered at baseline only.

### Behavioral outcomes

Four behavioral measures were analyzed across participant self-report and parent proxy-report; several reached nominal significance but none survived FDR correction, consistent with a seven-pair sample, so findings are interpreted as preliminary effect-size estimates. RCADS Total Anxiety, the primary outcome, decreased from pre to post with the maximal possible effect in both raters: self-report fell from 26.7 to 18.1 (Δ = −8.6; W = 0, p = 0.016; rrb = −1.00) and parent-report from 30.0 to 18.6 (Δ = −11.4; W = 0, p = 0.016; rrb = −1.00), every participant decreasing on both (Table 2, Figure 1). The reduction extended across subdomains: on self-report, Social Phobia decreased significantly (Δ = −3.9, p = 0.047), the remaining anxiety subscales and Total Internalizing showed large same-direction effects (Total Internalizing Δ = −11.9, rrb = −1.00), and a convergent pattern appeared on parent-report (Table 3). Across raters, every reported subscale moved toward reduced anxiety.

**Figure 1.**
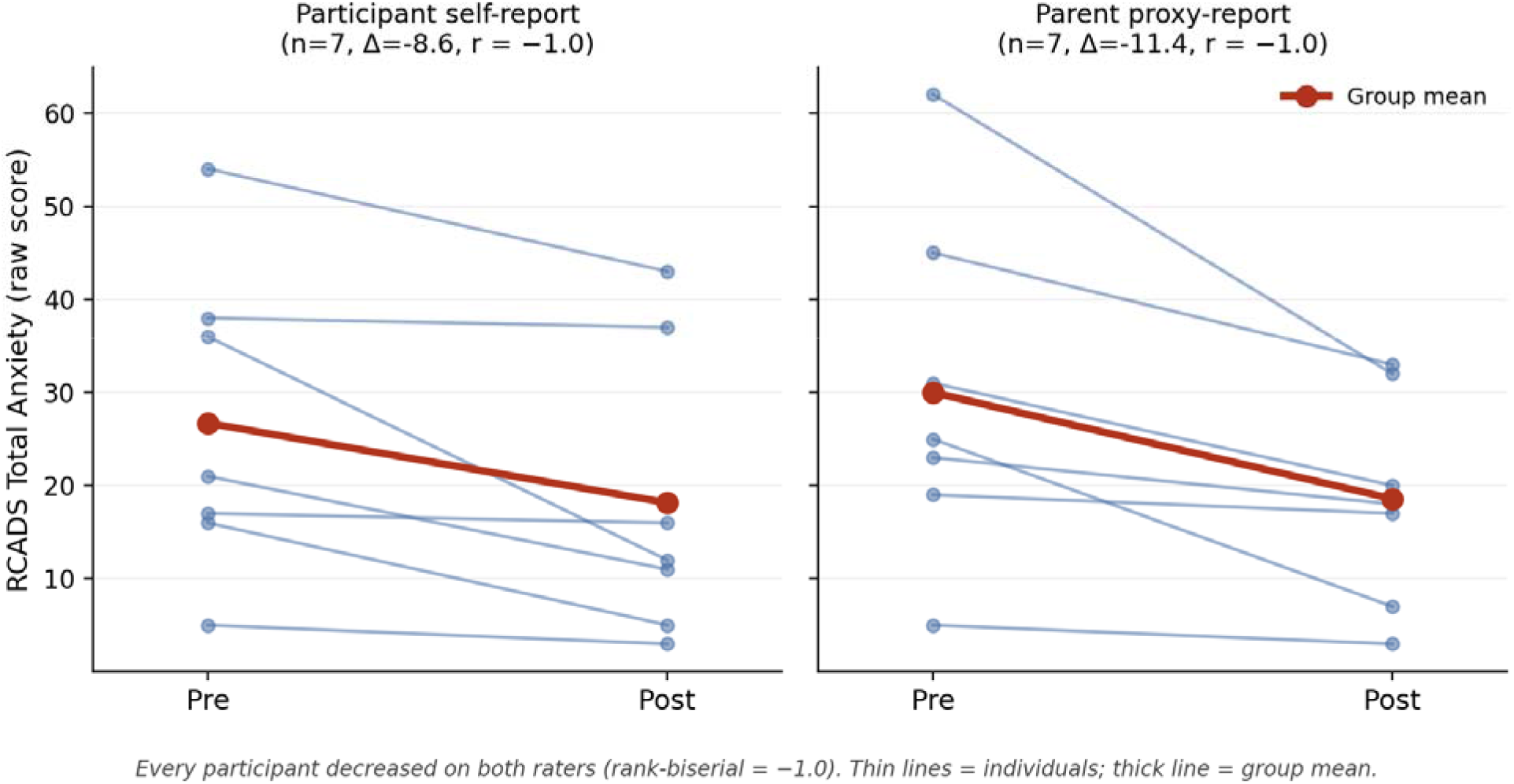
Individual participant trajectories in RCADS Total Anxiety from pre to post, shown separately for participant self-report and parent/guardian proxy-report. Each thin line is one participant or parent/guardian (n = 14); the heavy line is the group mean. All participants and parents/guardians decreased for both raters (rank-biserial = −1.00), though the magnitude of change varied, with two near-flat self-report cases (Δ = −1).

**Table 2.**
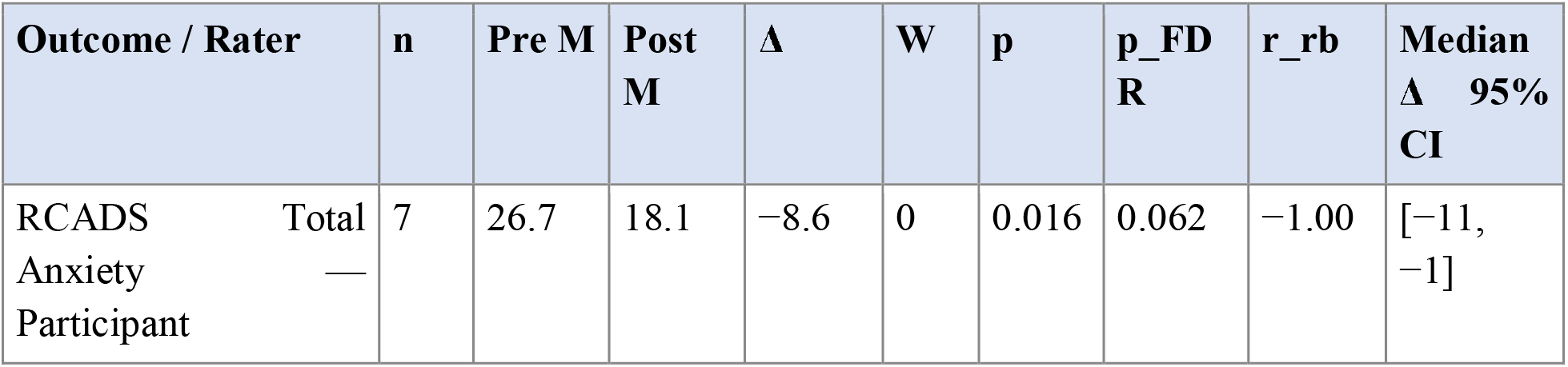
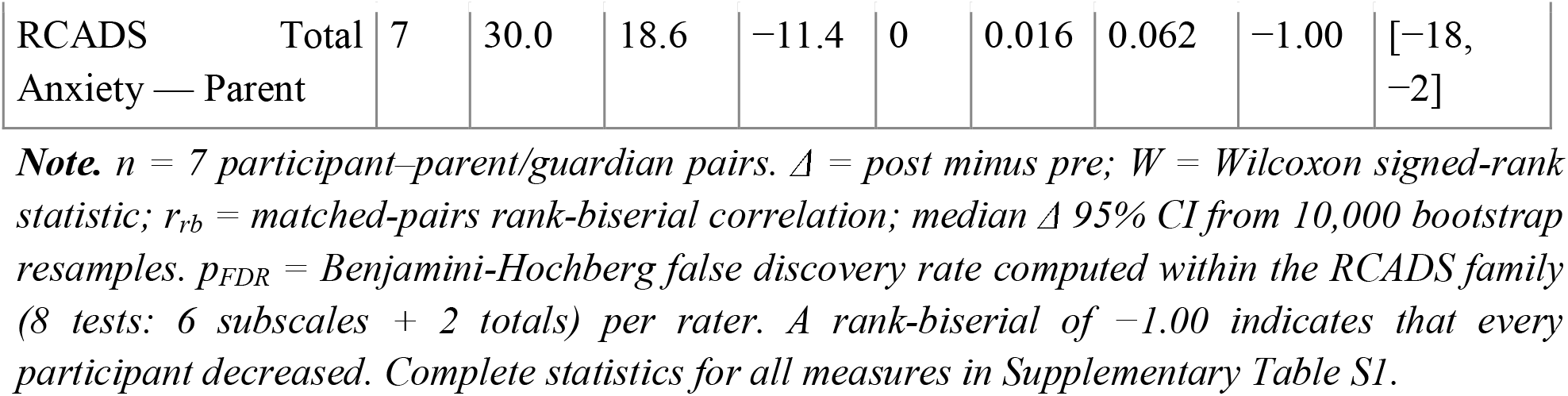
Pre-to-post change in the primary behavioral outcome (RCADS Total Anxiety), by rater.

**Table 3.**
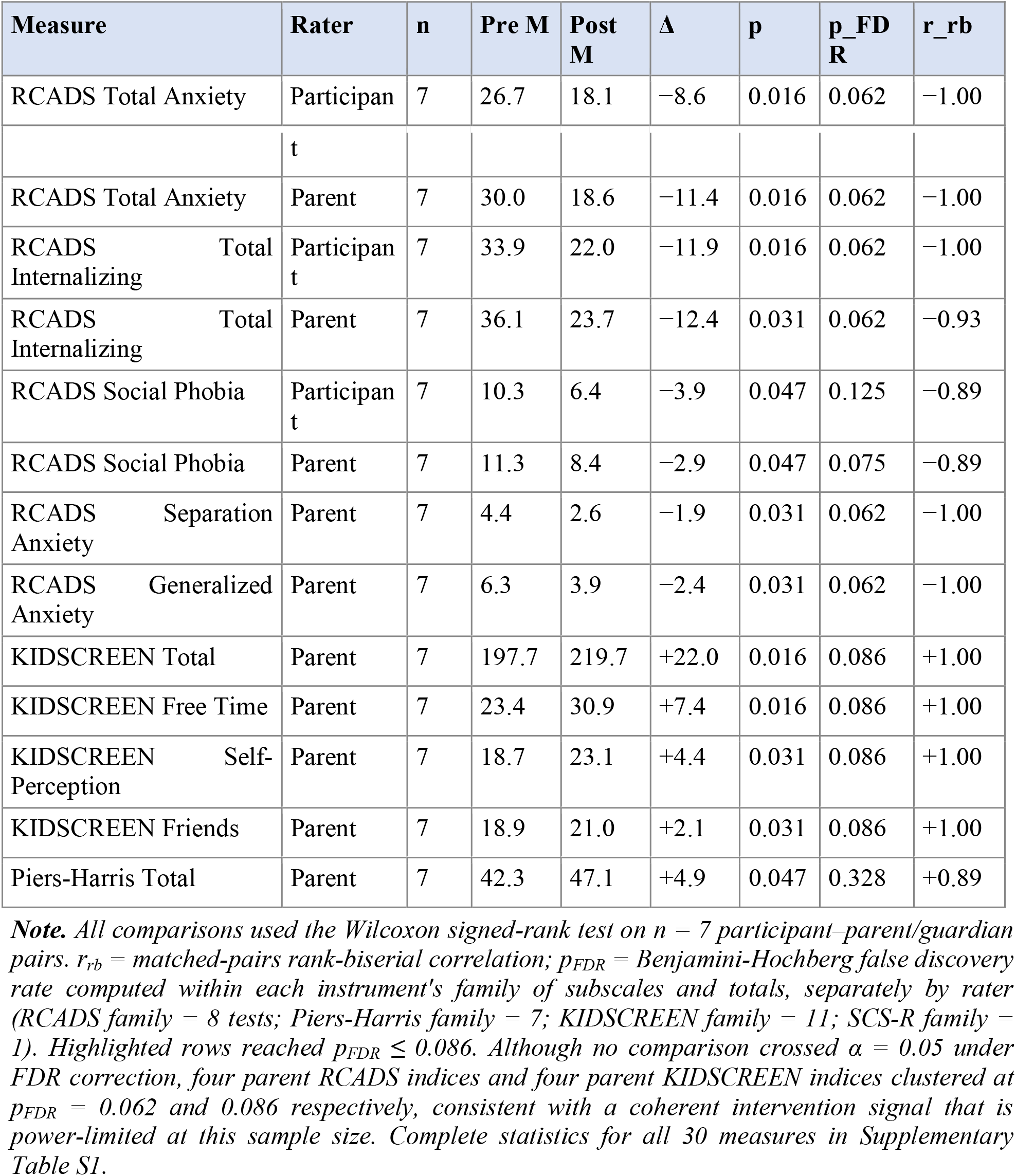
All behavioral outcomes reaching nominal significance (p < 0.05), ordered by instrument and rater.

| Measure | Rater | n | Pre M | Post M | $\Delta$ | p | $p_{FDR}$ | $r_{rb}$ |
| --- | --- | --- | --- | --- | --- | --- | --- | --- |
| RCADS Total Anxiety | Participan | 7 | 26.7 | 18.1 | -8.6 | 0.016 | 0.062 | -1.00 |

|  | t |  |  |  |  |  |  |  |
| --- | --- | --- | --- | --- | --- | --- | --- | --- |
| RCADS Total Anxiety | Parent | 7 | 30.0 | 18.6 | -11.4 | 0.016 | 0.062 | -1.00 |
| RCADS Total Internalizing | Participant | 7 | 33.9 | 22.0 | -11.9 | 0.016 | 0.062 | -1.00 |
| RCADS Total Internalizing | Parent | 7 | 36.1 | 23.7 | -12.4 | 0.031 | 0.062 | -0.93 |
| RCADS Social Phobia | Participant | 7 | 10.3 | 6.4 | -3.9 | 0.047 | 0.125 | -0.89 |
| RCADS Social Phobia | Parent | 7 | 11.3 | 8.4 | -2.9 | 0.047 | 0.075 | -0.89 |
| RCADS Separation Anxiety | Parent | 7 | 4.4 | 2.6 | -1.9 | 0.031 | 0.062 | -1.00 |
| RCADS Generalized Anxiety | Parent | 7 | 6.3 | 3.9 | -2.4 | 0.031 | 0.062 | -1.00 |
| KIDSCREEN Total | Parent | 7 | 197.7 | 219.7 | +22.0 | 0.016 | 0.086 | +1.00 |
| KIDSCREEN Free Time | Parent | 7 | 23.4 | 30.9 | +7.4 | 0.016 | 0.086 | +1.00 |
| KIDSCREEN Self-Perception | Parent | 7 | 18.7 | 23.1 | +4.4 | 0.031 | 0.086 | +1.00 |
| KIDSCREEN Friends | Parent | 7 | 18.9 | 21.0 | +2.1 | 0.031 | 0.086 | +1.00 |
| Piers-Harris Total | Parent | 7 | 42.3 | 47.1 | +4.9 | 0.047 | 0.328 | +0.89 |

**Note.** All comparisons used the Wilcoxon signed-rank test on $n = 7$ participant–parent/guardian pairs. $r_{rb}$ = matched-pairs rank-biserial correlation; $p_{FDR}$ = Benjamini-Hochberg false discovery rate computed within each instrument's family of subscales and totals, separately by rater (RCADS family = 8 tests; Piers-Harris family = 7; KIDSCREEN family = 11; SCS-R family = 1). Highlighted rows reached $p_{FDR} \leq 0.086$ . Although no comparison crossed $\alpha = 0.05$ under FDR correction, four parent RCADS indices and four parent KIDSCREEN indices clustered at $p_{FDR} = 0.062$ and $0.086$ respectively, consistent with a coherent intervention signal that is power-limited at this sample size. Complete statistics for all 30 measures in Supplementary Table S1.

On parent-report, secondary domains also improved: KIDSCREEN-52 Total rose (Δ = +22.0, p = 0.016, rrb = +1.00), as did Piers-Harris Total (Δ = +4.9, p = 0.047) and, at trend level, social connectedness (Δ = +8.0, p = 0.094); none survived within-instrument FDR correction. Participant self-report on these domains was largely unchanged (self-report KIDSCREEN Total Δ = −3.0, ns) — the most striking rater divergence in the data. Baseline agreement between raters was strong for the two measures with the largest effects (RCADS Total Anxiety and Piers-Harris Total, both ρ ≈ 0.80, p < 0.003), but change-score agreement was weaker and, for self-concept, divergent (Supplementary Results; Supplementary Tables S2–S3). A qualitative analysis of open-ended responses converged on themes of belonging, competence through challenge, purpose, and community-sustaining infrastructure for families (Supplementary Results).

### Spectral power (PSD)

Within the pre-specified fast-band family (27 cells: 3 conditions × 3 regions × 3 bands; max-statistic permutation FWER), theatrical performance produced large, family-wise-significant increases in high-beta and gamma power across temporal, parietal, and prefrontal cortex; eight cells survived correction (dz = 1.05–1.48, all pFWER < 0.05), the strongest being temporal high-beta (dz = 1.48, pFWER = 0.003) (Table 4; Figure 2). Movement mirroring produced a prefrontally dominant pattern with five surviving cells (dz = 1.03–1.27), localized to the left prefrontal pole (Fp1 > Fp2; Supplementary Figure S1). Paced breathing showed the opposite direction — a parietal and temporal gamma reduction (dz = −0.77 to −0.93) that did not survive correction but converged with the source-space connectivity results below.

**Figure 2.**
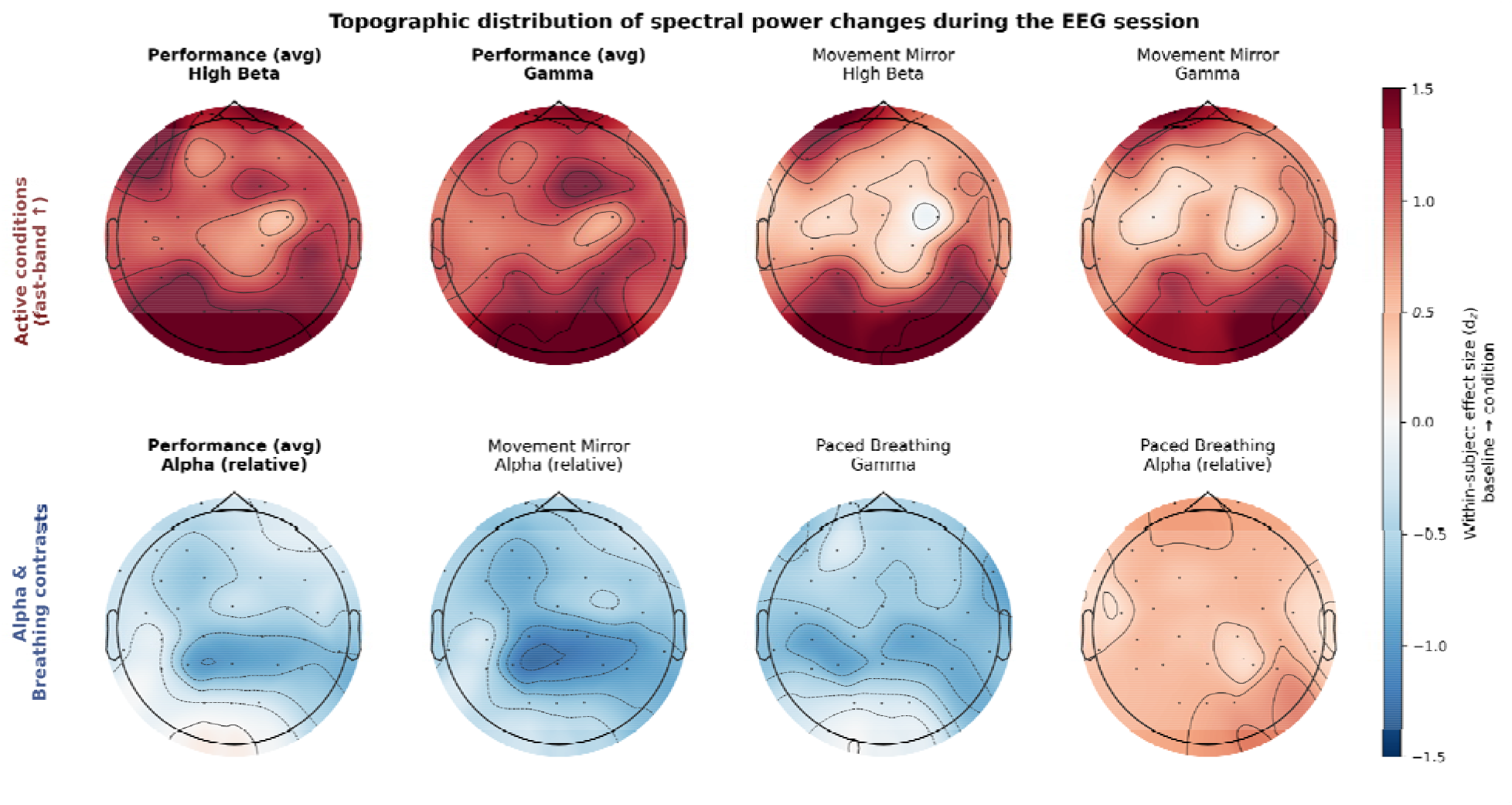
Topographic distribution of spectral power changes during the EEG session. Scalp maps show per-channel within-subject effect sizes (d_z_) for power change relative to resting baseline, rendered at the 32-channel LiveAmp electrode positions and interpolated across the scalp. Top row: During theatrical performance and movement mirroring, fast-band (high-beta and gamma) power increases broadly, with performance producing a more global activation pattern and mirror showing a frontal–prefrontal-dominant pattern. Bottom row, first three panels: Active conditions produce relative alpha suppression, strongest at parietal and central-parietal electrodes — the canonical EEG signature of attentional engagement. Bottom row, last two panels: Paced breathing inverts this pattern, with reduced gamma (blue) and increased alpha (red), consistent with reduced cortical engagement during the paced breathing condition. Positive d_z_ (red) indicates power increase relative to baseline; negative (blue) indicates decrease. N = 9–12 per condition. Per-channel d_z_ values are slightly smaller in magnitude than the ROI d_z_ reported in Table 4 (which standardize on ROI-averaged power, reducing within-subject noise before computing effect size); the spatial pattern is unchanged.

**Table 4.** Spectral power changes relative to resting baseline surviving family-wise correction by condition, region, and band.

| Condition vs. baseline | Region | Band | $d_z$ | $p_{FWER}$ |
| --- | --- | --- | --- | --- |
| Performance avg | Temporal | High beta | +1.48 | 0.003 |
| Performance avg | Prefrontal | Gamma | +1.43 | 0.005 |
| Performance avg | Prefrontal | High beta | +1.39 | 0.008 |
| Performance avg | Parietal | High beta | +1.29 | 0.013 |
| Performance avg | Parietal | Gamma | +1.22 | 0.022 |
| Performance avg | Temporal | Gamma | +1.20 | 0.022 |
| Performance avg | Temporal | Low beta | +1.06 | 0.043 |
| Performance avg | Prefrontal | Low beta | +1.05 | 0.045 |
| Mirror | Prefrontal | Gamma | +1.27 | 0.016 |
| Mirror | Prefrontal | Low beta | +1.24 | 0.017 |
| Mirror | Prefrontal | High beta | +1.17 | 0.029 |
| Mirror | Parietal | High beta | +1.17 | 0.029 |
| Mirror | Parietal | Gamma | +1.03 | 0.048 |

**Note.** $d_z$ = within-subject standardized effect size; positive = power increase, negative = power decrease. $p_{FWER}$ = family-wise error rate from max-statistic sign-flip permutation across the pre-specified 27-cell fast-band family (temporal, parietal, prefrontal × low beta, high beta, gamma). All 13 cells surviving family-wise correction ( $p_{FWER} < 0.05$ ) are shown; the complete 27-cell grid appears in Supplementary Table S4. $N = 12$ per cell.

The power effects were independent of age: all 13 survivors remained significant with age as a covariate (adjusted p ≤ 0.006), and no cell showed a significant age association (Supplementary Results). They were not fully independent of movement, however. The high-beta power increase scaled with concurrent movement vigor across participants (whole-head ρ = 0.86, LOO-robust), but spectral parameterization localized this coupling to the broadband (aperiodic) component: the oscillatory high-beta and gamma peaks were unrelated to movement (ρ = −0.06 and −0.22), so the fast-band effect comprises a movement-robust oscillatory component and a broadband component that co-occurs with movement and cannot, at this sample size, be separated from residual myogenic activity (Supplementary Results).

Outside the pre-specified family, an exploratory analysis of relative alpha power showed coherent parietal alpha suppression during every active condition (dz = −0.68 to −1.10), with the reverse during breathing; spectral parameterization indicated a genuine oscillatory-peak reduction that did not covary with movement (ρ = 0.08), consistent with attentional desynchronization (Figure 2; Supplementary Results).

### Intra-brain synchrony

All 78 recordings passed the rank screen (effective rank 19–30). The pre-specified high-frequency omnibus — each participant’s mean source-space MIM change across the 28 Yeo network pairs in high-beta and gamma — increased during performance relative to baseline (dz = 0.64, t(11) = 2.22, p = 0.049; Figure 3a). The effect was band-specific (high-beta dz = 0.61, gamma dz = 0.63; no increase in theta, alpha, or low-beta, all p > 0.21) and condition-specific (positive but non-significant during mirroring, dz = 0.41; absent during breathing, dz = −0.10; Supplementary Figure S2), and appeared in both within- and between-network coupling, marginally favoring between-network integration (Supplementary Figure S3).

**Figure 3.**
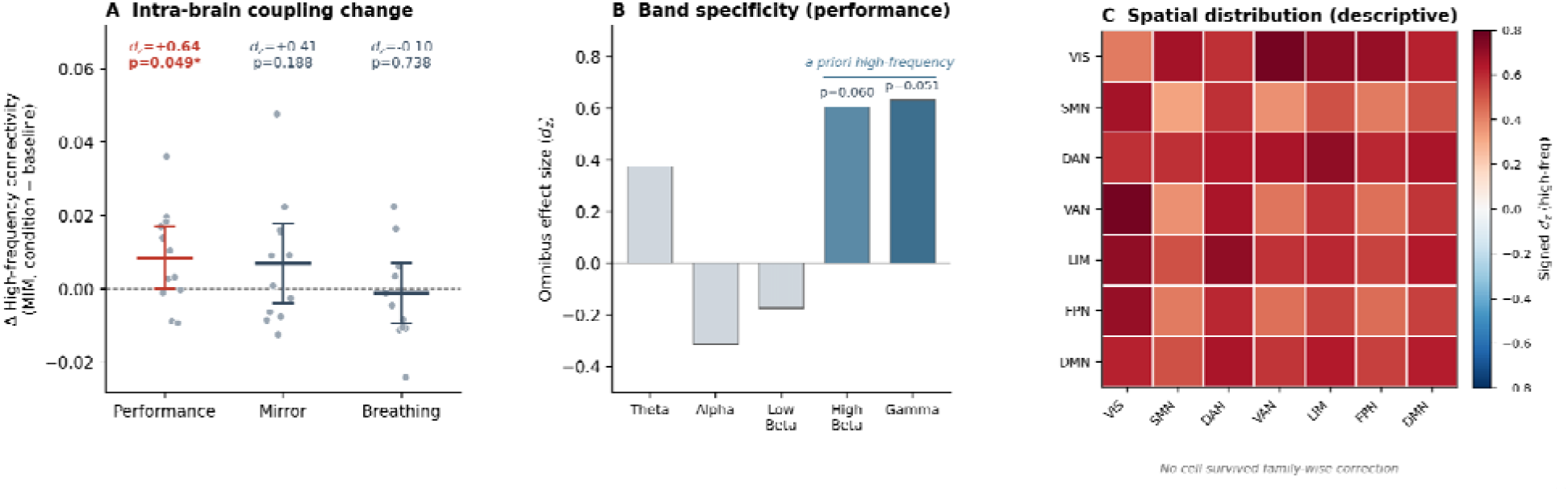
Intra-brain functional connectivity (source-space MIM, Yeo seven-network parcellation) during musical-theater engagement, relative to resting baseline. **(A)** Change in high-frequency (high-beta + gamma) intra-brain connectivity for each condition, expressed as the per-participant mean change across all 28 network pairs. Gray points are individual participants (n = 12); horizontal bars and whiskers denote the group mean and 95% confidence interval. Theatrical performance produced a significant overall increase in high-frequency coupling (d_z = 0.64, p = 0.049), whereas movement mirroring (d_z = 0.41) and paced breathing (d_z = −0.10) did not. **(B)** Band specificity of the performance effect: the omnibus connectivity increase was carried by the high-beta and gamma bands (highlighted) and was absent in theta, alpha, and low beta, matching the band localization of the spectral-power findings and the a priori high-frequency prediction. **(C)** Spatial distribution of the performance-related high-frequency change across the 7 × 7 network grid, shown as signed within-subject effect sizes (d_z; warm colors indicate increased coupling). The increase was broadly distributed and positive across all network pairs, with the largest descriptive effects in visual-, dorsal-attention–, and limbic-related couplings. No individual network pair survived family-wise correction (max-statistic sign-flip permutation across the full 28-pair × 5-band grid); panel C is therefore presented as a descriptive, hypothesis-generating depiction of where the distributed increase was strongest. MIM = multivariate interaction measure; d_z = within-subject standardized effect size.

The increase was carried by amplitude as well as phase: re-estimating the contrast with orthogonalized amplitude-envelope correlation yielded a much larger whole-brain effect (dz = 1.85, p < 0.001) than MIM (dz = 0.62) or absolute coherence (dz = 0.66), again concentrated in high-beta and gamma and graded across conditions (mirroring dz = 1.47; breathing dz = −0.53). Phase- and amplitude-based increases were only moderately correlated across participants (ρ = 0.57), indicating partially distinct contributions, and the connectivity change was not significantly associated with the concurrent power change (ρ = 0.38, p = 0.23). Two limits qualify the omnibus: under leave-one-out resampling its p ranged 0.024–0.094 (below 0.05 in 4 of 12 folds; Supplementary Figure S4), and no individual network pair survived correction, so the broadly distributed cell-level pattern is descriptive only (Supplementary Results).

The effect was independent of age (age-adjusted dz = 0.61; age slope p = 0.76) and of movement: both leakage-robust measures were uncorrelated with vigor (MIM _ρ_ = 0.43, p = 0.167; AEC _ρ_ = 0.04, p = 0.897), whereas the leakage-susceptible coherence did track vigor (_ρ_ = 0.62, p = 0.031) — a positive control confirming the accelerometry analysis detects movement when present (Supplementary Table S5). Convergently, amplitude coupling was comparably elevated during mirroring, a condition in which vigor did not rise above baseline.

### Inter-brain synchrony

To test whether shared performance produced coupling between as well as within brains, we compared real co-performing dyads against same-condition pseudo-dyads across five connectivity measures spanning phase (PLV, imaginary coherence), combined phase- and-amplitude (coherence), and amplitude (amplitude-envelope correlation; orthogonalized cross-brain AEC). This was a planned aim, but the analyses were not among the confirmatory, family-wise-corrected tests and rested on a few dyads (n = 7–10), so we report them as effect-size estimates. As within a single brain, the effect was carried by amplitude, not phase: no phase measure exceeded the pseudo-dyad surrogate in any number or focal band (PLV |d| ≤ 0.72; imaginary coherence largest d = 0.70), whereas the two amplitude measures distinguished real from pseudo-dyads (Figure 4; full grid in Supplementary Table S6).

**Figure 4.**
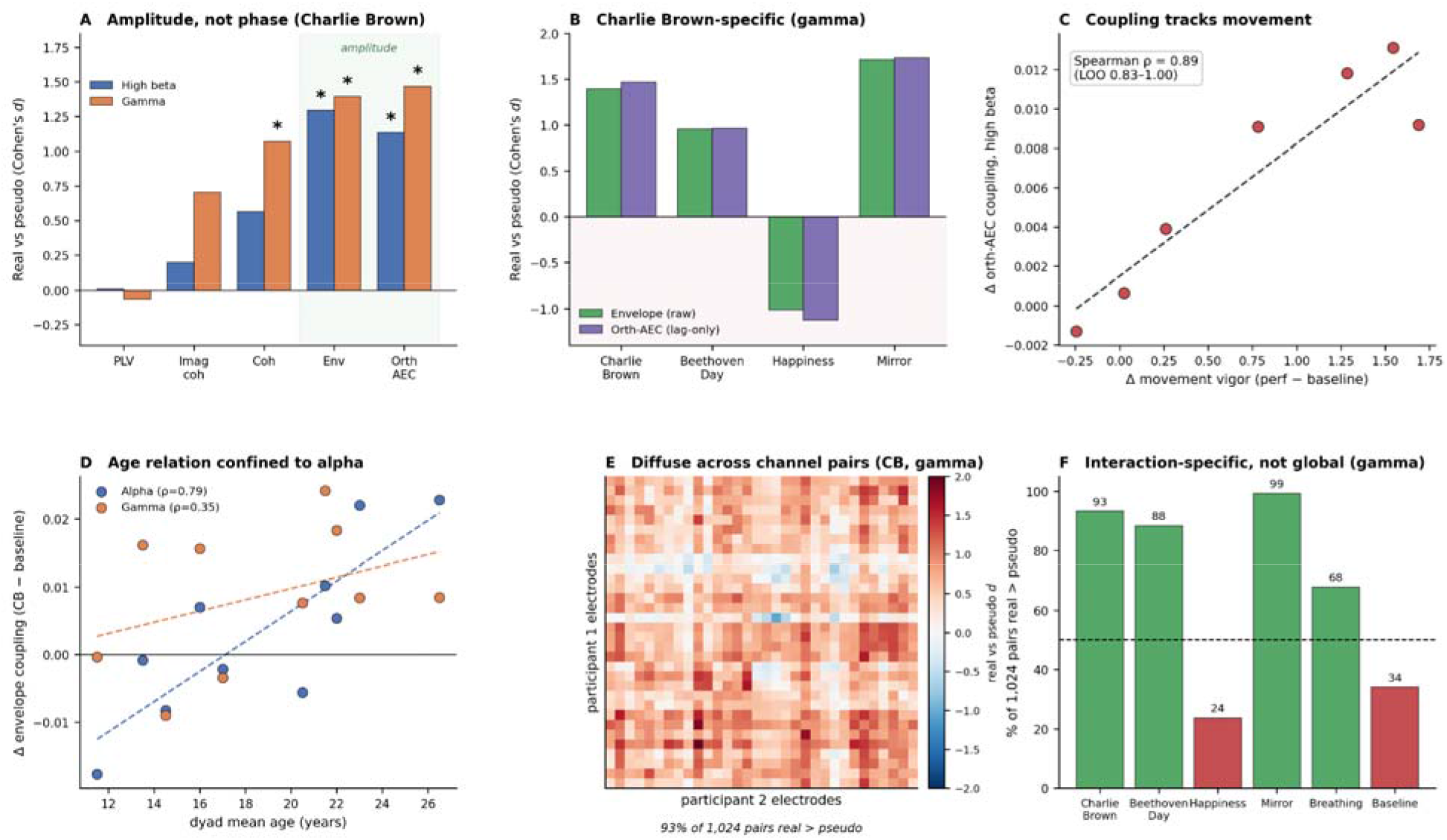
Inter-brain synchrony during shared performance. (A) Inter-brain coupling above the pseudo-dyad surrogate (Cohen’s d, real vs. pseudo) during the “You’re a Good Man, Charlie Brown” number for the five connectivity measures in the two focal bands; phase measures (PLV, imaginary coherence) do not exceed surrogate, whereas the amplitude measures (envelope correlation, orthogonalized cross-brain AEC) do, with coherence intermediate. Asterisks mark effects surviving Benjamini–Hochberg FDR. (B) Amplitude coupling above surrogate by condition (gamma), for the raw envelope correlation and the leakage-robust orthogonalized AEC; the two agree closely (indicating a genuine lagged component), and coupling is present for “You’re a Good Man, Charlie Brown,” “Beethoven Day,” and the mirror exercise (strongest effect) but absent (below surrogate) for the full-ensemble “Happiness” finale. (C) The performance-related increase in orthogonalized (lag-only) high-beta coupling scales across dyads with the concurrent increase in movement vigor (Spearman ρ = 0.89, leave-one-out 0.83–1.00; n = 7); the raw envelope measure shows the same relation (ρ = 0.86). (D) Dyad-level change in envelope coupling during “You’re a Good Man, Charlie Brown,” relative to baseline, as a function of dyad mean age: alpha coupling scales with age (ρ = 0.79) whereas gamma does not (ρ = 0.35), confining the age relation to alpha. (E) Spatial distribution: per-pair real-versus-surrogate coupling (Cohen’s d) across the full 32 × 32 cross-brain channel grid for “You’re a Good Man, Charlie Brown” (gamma); warm values fill the grid, and real coupling exceeds surrogate at 93% of the 1,024 pairs with no homologous-pair preference and no concentration in a small subset. (F) Proportion of the 1,024 cross-brain pairs at which real coupling exceeds surrogate, by condition (gamma); “You’re a Good Man, Charlie Brown,” “Beethoven Day,” and mirror exceed 50% (green), whereas the “Happiness” finale and resting baseline fall below (red). n = 7–10 dyads per condition. AEC = amplitude-envelope correlation; PLV = phase-locking value.

Among the three musical numbers, amplitude coupling was largest for “You’re a Good Man, Charlie Brown,” exceeding surrogate in both focal bands (high-beta d = 1.29, gamma d = 1.39; both q = 0.004; Table 5). The orthogonalized measure, which removes the instantaneously shared zero-lag component, retained roughly half the magnitude and remained above surrogate (high-beta d = 1.13, gamma d = 1.47), indicating a genuine lagged component not attributable to shared input; the same dyads also exceeded their own baseline (Table 5, Panel C). “Beethoven Day” showed the same direction subthreshold, and the “Happiness” finale showed no interaction-specific coupling. Extending the comparison to all conditions showed coupling was not specific to one number but graded with interpersonal movement coordination (Supplementary Table S7; Supplementary Figure S5): it was strongest during movement mirroring (envelope d = 1.59 and 1.71; orthogonalized d = 1.65 and 1.74; all p < 0.001), elevated during the dyadic numbers, moderate during breathing, and absent or below surrogate during the ensemble finale and rest — matching the coordination each condition demands. This selectivity contrasted with the broad, comparable increases in local power and within-brain connectivity across the three numbers (Supplementary Table S8).

**Table 5.** Inter-brain synchrony: real-versus pseudo-dyad comparisons, leakage-robust sensitivity analyses, and performance-versus-baseline change with age adjustment.

| Number | Band | PLV $d(q)$ | Coherence $d(q)$ | Envelope $d(q)$ |
| --- | --- | --- | --- | --- |
| You’re a Good Man, Charlie Brown | High beta | 0.01 (0.98) | 0.56 (0.20) | **1.29 (0.004)** |
| You’re a Good Man, Charlie Brown | Gamma | −0.07 (0.96) | **1.07 (0.02)** | **1.39 (0.004)** |
| Beethoven Day | High beta | 0.03 (0.98) | 0.39 (0.52) | 1.11 (0.07) |
| Beethoven Day | Gamma | 0.91 (0.11) | 1.01 (0.09) | 0.95 (0.10) |
| Happiness | High beta | −0.72 (0.20) | −0.59 (0.29) | −0.67 (0.22) |
| Happiness | Gamma | 0.29 (0.66) | 0.10 (0.96) | −1.01 (0.09) |

| Number | Band | Imaginary coh. $d(q)$ | Orthogonalized AEC $d(q)$ |
| --- | --- | --- | --- |
| You’re a Good Man, Charlie Brown | High beta | 0.20 (0.67) | **1.13 (0.02)** |
| You’re a Good Man, Charlie Brown | Gamma | 0.70 (0.10) | **1.47 (0.003)** |
| Beethoven Day | High beta | 0.25 (0.67) | 0.90 (0.10) |
| Beethoven Day | Gamma | 0.62 (0.28) | 0.96 (0.10) |
| Happiness | High beta | 0.15 (0.73) | −0.52 (0.37) |
| Happiness | Gamma | −0.22 (0.67) | −1.13 (0.06) |

| Measure / contrast | Band | $n$ | $dz(p)$ | $\rho$ age (p) | Age- | Adj. $dz$ |
| --- | --- | --- | --- | --- | --- | --- |

|  |  |  |  |  | **slope p** |  |
| --- | --- | --- | --- | --- | --- | --- |
| Envelope: Performance<br>avg – baseline | Alpha | 7 | 0.78 (0.09) | 0.57 (0.18) | 0.36 | 0.78 |
|  | High beta | 7 | 1.39 (0.02) | –0.54<br>(0.22) | 0.15 | 1.61 |
|  | Gamma | 7 | 1.23 (0.03) | –0.32<br>(0.48) | 0.47 | 1.19 |
| Envelope: You’re a<br>Good Man, Charlie<br>Brown – baseline | Alpha | 10 | 0.26 (0.44) | 0.79 (0.01) | 0.003 | 0.43 |
|  | High beta | 10 | 0.88 (0.03) | 0.33 (0.35) | 0.17 | 0.94 |
|  | Gamma | 10 | 0.83 (0.04) | 0.35 (0.33) | 0.28 | 0.84 |
| Orthog. AEC: Performance<br>avg – baseline | High beta | 7 | 1.19 (0.05) | –0.57<br>(0.18) | 0.26 | 1.37 |
|  | Gamma | 7 | 1.55 (0.02) | –0.25<br>(0.59) | 0.35 | 1.70 |
| Orthog. AEC: You’re a<br>Good Man, Charlie<br>Brown – baseline | High beta | 10 | 0.81 (0.04) | 0.31 (0.38) | 0.22 | 0.90 |
|  | Gamma | 10 | 0.96 (0.02) | 0.32 (0.36) | 0.27 | 1.04 |

The strength of coupling scaled with movement: across dyads, the performance-related increase in coupling tracked the concurrent increase in vigor, robustly in high-beta (envelope ρ = 0.86; orthogonalized ρ = 0.89; both LOO-stable), though more weakly within “You’re a Good Man, Charlie Brown” alone (ρ ≈ 0.3–0.4, ns). Because the association held for the orthogonalized measure, which removes the zero-lag component, the temporally lagged component of the coupling also co-varied with movement. This contrasts with the movement-independent within-brain amplitude coupling. The “You’re a Good Man, Charlie Brown” effect was robust to individual dyads and spatially diffuse, and the high-frequency coupling was age-independent (Supplementary Results; Supplementary Figure S6, Supplementary Table S9); estimates rest on few dyads (n = 7–10), so permutation p-values are approximate (dyad roster in Supplementary Table S10).

## DISCUSSION

Over a 16-week musical-theater program, adolescents and adults with developmental disabilities showed reduced anxiety together with a reorganization of neural activity that we resolved across three levels of description: spectral, within-brain, and between-brain. Three findings converged. Anxiety declined substantially from pre- to post-intervention, a reduction reported concordantly by participants and parents/guardians. During performance, fast-band (high-beta and gamma) power and within-brain amplitude connectivity rose broadly across the cortex, with maximal effects in parieto-occipital regions. Between-brain amplitude coupling was not general but graded by the degree of interpersonal movement coordination, peaking when partners most directly mirrored one another. All together, these results point to a single organizing principle; namely, that coordinated, socially embedded movement scaffolds neural activity both within and between brains — the central prediction of the Synchronicity Hypothesis of Dance(Basso *et al*., 2020). We interpret all estimates as effect sizes from a small, single-arm sample rather than as confirmatory tests.

The study’s central contribution is a dissociation in how shared performance reorganized activity across scales. Local fast-band power and within-brain connectivity rose comparably across all three musical numbers (Supplementary Figure S7), consistent with a general state of heightened arousal(Jensen *et al*., 2007), sensorimotor engagement(Engel and Fries, 2010), and movement common to any active performance(Muthukumaraswamy, 2013). Between-brain coupling behaved differently: it was selective — largest for the most interactive number and, across all conditions, strongest during the movement-mirroring exercise. Both the within- and between-brain connectivity increases were carried by amplitude co-fluctuation rather than phase locking, and the same amplitude-over-phase profile at both scales suggests that a single form of coupling, the coordinated waxing and waning of oscillatory strength, operated across levels. What differed was not the type of coupling but its breadth: this shared envelope-based mechanism produced a broad, state-like change within brains alongside a change between brains that was specific to the interpersonal structure of the task. That layering is the informative result. A broad within-brain change and a coordination-specific between-brain change are not the same phenomenon at two scales, but two dissociable signatures of joint performance.

Both the frequency and the form of the between-brain effect are interpretable, and both connect it to existing work. Amplitude-envelope coupling and phase coupling are distinct intrinsic coupling modes with different origins and putative functions(Hipp *et al*., 2012): phase alignment is the substrate of precise, cycle-by-cycle neuronal communication(Fries, 2015), whereas envelope co-fluctuation indexes the co-modulation of oscillatory strength (i.e., the shared engagement of two systems ramping up and down together). That the coupling was envelope-based rather than phase-locked therefore points to coordinated engagement rather than millisecond-precise information exchange, an interpretation more plausible across two separate skulls, where consistent gamma phase alignment without a shared driver is biophysically unlikely. The high-frequency locus fits the same reading. Beta-band inter-brain coupling is consistently reported in social settings involving joint action, action observation, and imitation, generally interpreted as sensorimotor, since beta reflects motor preparation and the maintenance of the current sensorimotor state(Engel and Fries, 2010). For a task built on coordinated movement, high-beta coupling is exactly where an interpersonally coordinated sensorimotor signal should appear. Gamma inter-brain coupling has previously been observed during spontaneous imitation, alongside alpha-mu and beta(Dumas *et al*., 2010), and linked to prosocial coordination — though it remains less examined and more debated, partly because its fast, transient dynamics resist a simple mapping onto slow behavioral coordination and partly because many hyperscanning studies restrict analysis to lower frequencies. By retaining and source-localizing the full high-frequency range with leakage-robust amplitude measures, we characterize coupling that much of the field does not examine, and its convergence with movement coordination suggests that the shared, coordinated sensorimotor engagement of joint performance, not abstract social inference, is what co-modulates the two brains.

The between-brain gradient is the study’s most direct test of the Synchronicity Hypothesis, and three features align with it: the coupling was amplitude-based, it was strongest where interpersonal movement coordination was greatest, and its magnitude scaled across dyads with the concurrent increase in movement vigor. This movement-dependence is precisely what the hypothesis predicts: coordinated movement is proposed to drive between-brain synchrony, and it is what most clearly distinguishes the between-brain effect from the movement-independent within-brain changes. The same property makes the effect harder to interpret, because coordinated movement can generate lagged amplitude coupling of non-neural origin, and the coupling peaked during mirroring, when partners’ movements were most correlated. Our design constrains this alternative: the pseudo-dyad control removes coupling attributable to shared task structure, and the orthogonalized measure, which strips out the instantaneously shared zero-lag component, still reproduced the full gradient. Therefore, the effect is not a trivial consequence of simultaneous motion. What neither control removes is temporally lagged movement coordination, and disentangling movement-driven neural synchrony from lagged movement coordination per se is the pivotal question these data raise — a tractable one, given that the gradient is already robust to the leakage-based controls available here.

Positioned against the existing literature, several features of this work are, to our knowledge, new. Hyperscanning in ASD and related developmental disabilities remains small and has relied largely on functional near-infrared spectroscopy, phase-based measures, and child–caregiver or participant–experimenter dyads in conversation or simple imitation(Kruppa *et al*., 2020). The present study instead used EEG, which affords the oscillatory resolution such measures lack; examined peer co-performers engaged in a genuine theatrical production rather than experimenter-led dyads; quantified inter-brain coupling with source-localized, leakage-robust amplitude measures across the full high-frequency range; and did so under a graded movement-coordination manipulation with a pseudo-dyad control. The pattern is also notable against the modal finding that inter-brain synchrony is reduced in ASD; the same fNIRS study found no elevated synchrony in its ASD group(Kruppa *et al*., 2020); whereas here, a structured, coordinated, movement-rich activity elicited robust and systematically graded inter-brain synchrony in a predominantly ASD sample. Because we included no typically developing comparison group, this speaks to the presence and gradation of coupling under these conditions, not to its restoration; establishing whether coordinated performance recovers synchrony that is otherwise diminished is a question for a controlled design.

The convergent anxiety reduction, though from a single-arm study, is consistent with features that musical theater uniquely combines: structured rhythmic and coordinated movement, repeated performance within a supportive group, and sustained engagement toward a shared creative goal. Parents/guardians additionally reported gains across broader psychosocial domains that participants did not themselves report; such rater divergence is common and may reflect genuinely different vantage points or a response shift in self-appraisal over the program(Sprangers and Schwartz, 1999), and it cautions against treating any single instrument as definitive. Because the design lacked a control group, these improvements cannot be attributed specifically to the intervention rather than to time, repeated testing, or nonspecific attention; they motivate, but do not establish, efficacy. The contribution of musical theater here is as an ecologically meaningful vehicle for the coordinated movement the neural findings implicate, and as a setting in which multi-scale neural measurement during genuine joint action is feasible in this population.

These findings should be read as bounding the scope of inference, not as undermining the effects themselves. The sample was small (12 with usable EEG; seven behavioral completers) and diagnostically mixed — predominantly ASD, but including individuals with intellectual disability or a chromosomal condition — so the neural findings characterize a developmental-disability sample rather than an ASD-specific one. A typically developing comparison will be needed to establish specificity. The single-arm, pre–post design precludes causal attribution of the behavioral change. Our a-priori neural prediction concerned alpha, so the high-beta and gamma effects, though convergent across scales, are best treated as hypothesis-generating with respect to frequency. Additionally, the high-frequency omnibus effects were modest and, under leave-one-out resampling, sensitive to individual participants, as expected for a pilot of this size. What the design nonetheless establishes is a coherent, mechanistically specified target: an amplitude-based, high-frequency inter-brain coupling that scales with interpersonal movement coordination, survives leakage-robust control, and dissociates from the movement-independent within-brain changes. That specificity is what makes the confirmatory study tractable. A controlled trial with a comparison activity, motion capture to partial out movement, pre-registered high-frequency hypotheses, and a larger, more homogeneous dyad sample would directly test whether coordinated movement drives between-brain synchrony.

In conclusion, this pilot provides a proof of concept that a naturalistic musical-theater program can be studied at multiple neural scales at once, and yields a coherent, testable picture: broad, state-like changes in local and within-brain activity during performance, a between-brain coupling that tracks interpersonal movement coordination, and a convergent reduction in anxiety. The results are promising rather than definitive, and their central claim, that coordinated movement organizes multi-scale neural synchrony, remains to be separated from a movement-coordination confound. If the coordination gradient survives that test, musical theater would offer a rare combination: an intervention that is genuinely engaging for ASD participants and, at the same time, a tractable model for how shared movement shapes the social brain.

## Supporting information

Supplementary Methods and Results

## Author Contributions

J.C.B. designed the study. N.T. and J.C.B. collected the data. Z.M., N.T., and J.C.B. developed the analyses, wrote the analysis code, and analyzed the behavioral and neural data. J.C.B. advised analyses. Z.M., N.T., R.F., C.J., V.T., G.N., S.D., and J.C.B wrote the manuscript and all authors edited the manuscript.

## Acknowledgements

We are deeply grateful to the participants and their families, whose enthusiasm and trust made this work possible, and to the directors and staff of the STEP-VA program for their partnership throughout the study.

