## Supplementary Tables for "The Choreography of Coupled Brains: Coordinated Movement and Multi-Scale Neural Synchrony in Musical Theater for Individuals with Developmental Disabilities"

**Supplementary Table S1**

*Complete pre-to-post statistics for all behavioral measures, by rater.*

| Measure | Rater | n | Pre M | Post M | Δ | 95% CI (Δ med) | W | p | p_FDR | r_rb |
|---|---|---|---|---|---|---|---|---|---|---|
| RCADS Social Phobia | Self | 7 | 10.29 | 6.43 | -3.86 | [-8, -1] | 2 | .047 | .125 | -0.89 |
| RCADS Social Phobia | Parent | 7 | 11.29 | 8.43 | -2.86 | [-5, -1] | 2 | .047 | .075 | -0.89 |
| RCADS Panic Disorder | Self | 7 | 3.29 | 3.00 | -0.29 | [-1, +1] | 10 | 1.000 | 1.000 | -0.05 |
| RCADS Panic Disorder | Parent | 7 | 4.14 | 1.71 | -2.43 | [-4, +0] | 2 | .188 | .214 | -0.80 |
| RCADS Separation Anxiety | Self | 7 | 4.29 | 2.57 | -1.71 | [-3, +0] | 2 | .094 | .150 | -0.86 |
| RCADS Separation Anxiety | Parent | 7 | 4.43 | 2.57 | -1.86 | [-3, -1] | 0 | .031 | .063 | -1.00 |
| RCADS Generalized Anxiety | Self | 7 | 4.43 | 3.14 | -1.29 | [-4, +0] | 1 | .250 | .286 | -0.80 |
| RCADS Generalized Anxiety | Parent | 7 | 6.29 | 3.86 | -2.43 | [-4, -1] | 0 | .031 | .063 | -1.00 |
| RCADS OCD | Self | 7 | 4.43 | 3.00 | -1.43 | [-3, +0] | 3 | .156 | .208 | -0.71 |
| RCADS OCD | Parent | 7 | 3.86 | 2.00 | -1.86 | [-3, +0] | 0 | .063 | .083 | -1.00 |
| RCADS Major Depression | Self | 7 | 7.14 | 3.86 | -3.29 | [-5, +0] | 2 | .094 | .150 | -0.81 |
| RCADS Major Depression | Parent | 7 | 6.14 | 5.14 | -1.00 | [-3, -1] | 7 | .281 | .281 | -0.50 |
| RCADS Total Anxiety | Self | 7 | 26.71 | 18.14 | -8.57 | [-11, -1] | 0 | .016 | .063 | -1.00 |
| RCADS Total Anxiety | Parent | 7 | 30.00 | 18.57 | -11.43 | [-18, -2] | 0 | .016 | .063 | -1.00 |
| RCADS Total Internalizing | Self | 7 | 33.86 | 22.00 | -11.86 | [-22, -5] | 0 | .016 | .063 | -1.00 |
| RCADS Total Internalizing | Parent | 7 | 36.14 | 23.71 | -12.43 | [-19, -3] | 1 | .031 | .063 | -0.93 |
| Piers-Harris Behavioral Adjustment | Self | 7 | 8.43 | 8.86 | +0.43 | [+0, +1] | 0 | .500 | .700 | +1.00 |
| Piers-Harris Behavioral Adjustment | Parent | 7 | 9.00 | 9.57 | +0.57 | [+0, +1] | 0 | .500 | .583 | +1.00 |
| Piers-Harris Freedom from Anxiety | Self | 7 | 5.57 | 6.29 | +0.71 | [+0, +1] | 0 | .063 | .438 | +1.00 |
| Piers-Harris Freedom from Anxiety | Parent | 7 | 4.57 | 5.57 | +1.00 | [+0, +3] | 2 | .250 | .350 | +0.73 |
| Piers-Harris Happiness/Satisfaction | Self | 7 | 8.71 | 9.14 | +0.43 | [+1, +1] | 7 | .359 | .700 | +0.50 |
| Piers-Harris Happiness/Satisfaction | Parent | 7 | 8.57 | 9.43 | +0.86 | [+0, +2] | 1 | .250 | .350 | +0.80 |
| Piers-Harris Intellectual/School | Self | 7 | 8.43 | 9.14 | +0.71 | [-2, +3] | 6 | .688 | .802 | +0.27 |
| Piers-Harris Intellectual/School | Parent | 7 | 8.14 | 9.00 | +0.86 | [+0, +2] | 3 | .156 | .350 | +0.76 |

| | | | | | | | | | | |
|---|---|---|---|---|---|---|---|---|---|---|
| Piers-Harris Physical Appearance | Self | 7 | 5.29 | 4.71 | -0.57 | [-1, +0] | 0 | .500 | .700 | -1.00 |
| Piers-Harris Physical Appearance | Parent | 7 | 5.00 | 5.29 | +0.29 | [+0, +1] | 2 | .750 | .750 | +0.50 |
| Piers-Harris Popularity | Self | 7 | 8.43 | 8.57 | +0.14 | [-1, +0] | 4 | 1.000 | 1.000 | -0.20 |
| Piers-Harris Popularity | Parent | 7 | 7.00 | 8.29 | +1.29 | [-1, +3] | 3 | .125 | .350 | +0.71 |
| Piers-Harris Total | Self | 7 | 45.29 | 46.71 | +1.43 | [-2, +6] | 10 | .484 | .700 | +0.32 |
| Piers-Harris Total | Parent | 7 | 42.29 | 47.14 | +4.86 | [+1, +10] | 2 | .047 | .328 | +0.89 |
| KIDSCREEN Health | Self | 7 | 17.86 | 18.00 | +0.14 | [-1, +3] | 9 | .422 | 1.000 | +0.39 |
| KIDSCREEN Health | Parent | 7 | 16.71 | 17.86 | +1.14 | [-1, +2] | 4 | .219 | .344 | +0.62 |
| KIDSCREEN Feelings | Self | 7 | 26.43 | 24.71 | -1.71 | [-5, +2] | 4 | .750 | 1.000 | -0.30 |
| KIDSCREEN Feelings | Parent | 7 | 22.43 | 25.29 | +2.86 | [+0, +5] | 0 | .063 | .115 | +1.00 |
| KIDSCREEN Mood | Self | 7 | 27.14 | 26.43 | -0.71 | [-1, +1] | 7 | 1.000 | 1.000 | +0.07 |
| KIDSCREEN Mood | Parent | 7 | 25.29 | 25.86 | +0.57 | [-2, +2] | 7 | .688 | .825 | +0.33 |
| KIDSCREEN Self-Perception | Self | 7 | 23.29 | 26.00 | +2.71 | [-2, +6] | 7 | .500 | 1.000 | +0.38 |
| KIDSCREEN Self-Perception | Parent | 7 | 18.71 | 23.14 | +4.43 | [+2, +8] | 0 | .031 | .086 | +1.00 |
| KIDSCREEN Free Time | Self | 7 | 25.14 | 25.43 | +0.29 | [-2, +4] | 7 | .875 | 1.000 | +0.13 |
| KIDSCREEN Free Time | Parent | 7 | 23.43 | 30.86 | +7.43 | [+4, +10] | 0 | .016 | .086 | +1.00 |
| KIDSCREEN Family | Self | 7 | 29.57 | 26.71 | -2.86 | [-4, +2] | 5 | .344 | 1.000 | -0.52 |
| KIDSCREEN Family | Parent | 7 | 28.57 | 28.86 | +0.29 | [-2, +4] | 10 | .906 | .906 | +0.10 |
| KIDSCREEN Financial | Self | 7 | 21.00 | 20.14 | -0.86 | [-2, +1] | 3 | .625 | 1.000 | -0.40 |
| KIDSCREEN Financial | Parent | 7 | 19.00 | 21.29 | +2.29 | [+0, +4] | 0 | .063 | .115 | +1.00 |
| KIDSCREEN Friends | Self | 7 | 20.86 | 19.86 | -1.00 | [-8, +5] | 9 | .719 | 1.000 | -0.19 |
| KIDSCREEN Friends | Parent | 7 | 18.86 | 21.00 | +2.14 | [+1, +4] | 0 | .031 | .086 | +1.00 |
| KIDSCREEN School | Self | 7 | 12.86 | 12.29 | -0.57 | [-2, +1] | 4 | .750 | 1.000 | -0.30 |
| KIDSCREEN School | Parent | 7 | 12.43 | 12.86 | +0.43 | [+0, +1] | 2 | .750 | .825 | +0.50 |
| KIDSCREEN Social Acceptance | Self | 7 | 12.57 | 14.14 | +1.57 | [+0, +3] | 0 | .250 | 1.000 | +1.00 |
| KIDSCREEN Social Acceptance | Parent | 7 | 12.29 | 12.71 | +0.43 | [+0, +1] | 0 | .500 | .688 | +1.00 |
| KIDSCREEN Total | Self | 7 | 216.71 | 213.71 | -3.00 | [-31, +17] | 13 | .938 | 1.000 | +0.07 |
| KIDSCREEN Total | Parent | 7 | 197.71 | 219.71 | +22.00 | [+10, +34] | 0 | .016 | .086 | +1.00 |
| SCS-R | Self | 7 | 95.29 | 99.71 | +4.43 | [-1, +6] | 6 | .344 | .344 | +0.48 |
| SCS-R | Parent | 7 | 82.86 | 90.86 | +8.00 | [+0, +18] | 2 | .094 | .094 | +0.86 |

***Note.*** *n = 7 complete pre/post pairs for all comparisons. Self = participant self-report; Parent = guardian proxy-report on the participant. Δ = post minus pre; W = Wilcoxon signed-rank statistic;* $r_{rb}$ *= matched-pairs rank-biserial correlation; 95% CI of median Δ from 10,000 bootstrap resamples.* $p_{FDR}$ *= Benjamini-Hochberg false discovery rate computed within each instrument's family (RCADS: 8 tests = 6 subscales + Total Anxiety + Total Internalizing; Piers-Harris: 7 tests = 6 subscales + Total; KIDSCREEN: 11 tests = 10 subscales + Total; SCS-R: 1 test), separately by rater. Highlighted rows have* $p_{FDR} < .10$*. KIDSCREEN parent Total was computed by summing the 10 parent subscale scores (no pre-computed total column was available in the source data; all 7 parents completed all 10 subscales). RCADS Total Anxiety = sum of the five anxiety subscales (Social Phobia, Panic Disorder, Separation Anxiety, Generalized Anxiety, OCD); RCADS Total Internalizing = Total Anxiety + Major Depression. PANAS and ICS measures were collected at pre and post but are not analyzed in this manuscript and are not shown.*

**Supplementary Table S2. Baseline ERS-ACA Subscales as Predictors of Outcome Change**

Spearman correlations between baseline ERS-ACA subscale scores and pre-to-post change in each outcome (n = 6; change scores sign-flipped so that positive values denote improvement). Leave-one-out (LOO) range gives the minimum and maximum ρ across single-case deletions. All analyses are exploratory; no FDR correction was applied. At n = 6 these estimates are unstable and are reported as hypothesis-generating only.

| ERS Subscale | Outcome (Δ, + = improvement) | ρ | p | n | LOO range |
|---|---|---|---|---|---|
| General Factor | KIDSCREEN Total | +0.93 | 0.008 | 6 | [+0.87, +0.97] |
| General Factor | RCADS Anxiety | -0.72 | 0.109 | 6 | [-0.95, -0.53] |
| General Factor | SCS | +0.64 | 0.173 | 6 | [+0.36, +0.97] |
| General Factor | PANAS Pos. Affect | +0.64 | 0.173 | 6 | [+0.36, +0.97] |
| General Factor | PANAS Neg. Affect | -0.18 | 0.738 | 6 | [-0.87, +0.05] |
| Approach Strategies | KIDSCREEN Total | +0.90 | 0.015 | 6 | [+0.82, +1.00] |
| Approach Strategies | RCADS Anxiety | -0.58 | 0.225 | 6 | [-0.89, -0.29] |
| Approach Strategies | SCS | +0.55 | 0.257 | 6 | [+0.21, +0.82] |
| Approach Strategies | PANAS Pos. Affect | +0.75 | 0.084 | 6 | [+0.56, +1.00] |
| Approach Strategies | PANAS Neg. Affect | -0.09 | 0.868 | 6 | [-0.72, +0.21] |
| Self-Development | KIDSCREEN Total | +0.88 | 0.021 | 6 | [+0.78, +0.89] |
| Self-Development | RCADS Anxiety | -0.38 | 0.464 | 6 | [-0.97, -0.16] |
| Self-Development | SCS | +0.39 | 0.439 | 6 | [+0.15, +0.67] |
| Self-Development | PANAS Pos. Affect | +0.52 | 0.295 | 6 | [+0.11, +0.89] |
| Self-Development | PANAS Neg. Affect | -0.03 | 0.954 | 6 | [-0.67, +0.29] |
| Avoidance Strategies | KIDSCREEN Total | +0.46 | 0.354 | 6 | [+0.10, +0.67] |
| Avoidance Strategies | RCADS Anxiety | -0.81 | 0.053 | 6 | [-0.97, -0.65] |
| Avoidance Strategies | SCS | +0.81 | 0.050 | 6 | [+0.67, +0.97] |
| Avoidance Strategies | PANAS Pos. Affect | +0.43 | 0.389 | 6 | [+0.10, +0.72] |
| Avoidance Strategies | PANAS Neg. Affect | -0.69 | 0.128 | 6 | [-0.76, -0.60] |

***Note.*** KIDSCREEN Total change was the only outcome with consistent large associations (General Factor ρ = .93; Approach ρ = .90; Self-Development ρ = .88). The Avoidance–SCS and Avoidance–RCADS associations reached nominal thresholds but are based on n = 6 and are not interpreted.

**Supplementary Table S3. Participant–Parent Agreement**

Spearman rank correlations between participant self-report and parent proxy-report, for baseline (pre) scores and for pre-to-post change scores. Pre-score agreement uses all dyads with baseline data for both raters (n = 12); change-score agreement uses paired completers (n = 7). Leave-one-out (LOO) range gives the minimum and maximum ρ across single-case deletions, indexing stability.

| **Measure** | **Type** | **ρ** | **p** | **n** | **LOO range** |
|---|---|---|---|---|---|
| RCADS Total Anxiety | pre | +.80 | .002 | 12 | [+.73, +.86] |
| RCADS Total Anxiety | change | +.56 | .191 | 7 | [+.37, +.79] |
| Piers-Harris Total | pre | +.78 | .003 | 12 | [+.71, +.93] |
| Piers-Harris Total | change | −.21 | .645 | 7 | [−.60, +.26] |
| SCS | pre | +.56 | .056 | 12 | [+.48, +.74] |
| SCS | change | +.61 | .148 | 7 | [+.43, +.77] |
| PANAS Positive | pre | +.32 | .306 | 12 | [+.17, +.44] |
| PANAS Positive | change | +.18 | .702 | 7 | [−.31, +.54] |
| PANAS Negative | pre | +.48 | .118 | 12 | [+.32, +.73] |
| PANAS Negative | change | +.01 | .985 | 7 | [−.60, +.28] |
| ICS Community | pre | +.13 | .692 | 12 | [−.10, +.32] |
| ICS Community | change | +.33 | .467 | 7 | [+.00, +.68] |
| ICS STEP-VA | pre | +.27 | .387 | 12 | [+.16, +.61] |
| ICS STEP-VA | change | +.64 | .119 | 7 | [+.42, +.85] |

*Note.* Correlations are descriptive; at n = 7 the change-score estimates are unstable, as the LOO ranges show. No correction for multiple comparisons was applied.

**Supplementary Table S4**

*Complete pre-registered fast-band family: within-subject spectral power change from resting baseline for all 27 cells (3 conditions × 3 regions × 3 bands).*

| Condition vs. baseline | Region | Band | $d_z$ | $p_{FWER}$ | Survived FWER |
|---|---|---|---|---|---|
| Performance avg | Temporal | High beta | +1.48 | 0.003 | Yes |
| Performance avg | Prefrontal | Gamma | +1.43 | 0.005 | Yes |
| Performance avg | Prefrontal | High beta | +1.39 | 0.008 | Yes |
| Performance avg | Parietal | High beta | +1.29 | 0.013 | Yes |
| Performance avg | Parietal | Gamma | +1.22 | 0.022 | Yes |
| Performance avg | Temporal | Gamma | +1.20 | 0.022 | Yes |
| Performance avg | Temporal | Low beta | +1.06 | 0.043 | Yes |
| Performance avg | Prefrontal | Low beta | +1.05 | 0.045 | Yes |
| Performance avg | Parietal | Low beta | +0.87 | 0.120 | — |
| Mirror | Prefrontal | Gamma | +1.27 | 0.016 | Yes |
| Mirror | Prefrontal | Low beta | +1.24 | 0.017 | Yes |
| Mirror | Parietal | High beta | +1.17 | 0.029 | Yes |
| Mirror | Prefrontal | High beta | +1.17 | 0.029 | Yes |
| Mirror | Parietal | Gamma | +1.03 | 0.048 | Yes |
| Mirror | Temporal | High beta | +0.87 | 0.118 | — |
| Mirror | Temporal | Low beta | +0.76 | 0.203 | — |
| Mirror | Temporal | Gamma | +0.75 | 0.214 | — |
| Mirror | Parietal | Low beta | +0.70 | 0.255 | — |
| Breathing | Parietal | Gamma | −0.93 | 0.085 | — |
| Breathing | Temporal | Gamma | −0.77 | 0.197 | — |
| Breathing | Temporal | Low beta | −0.48 | 0.625 | — |
| Breathing | Parietal | Low beta | −0.48 | 0.628 | — |
| Breathing | Parietal | High beta | −0.47 | 0.637 | — |
| Breathing | Temporal | High beta | −0.47 | 0.642 | — |
| Breathing | Prefrontal | Gamma | −0.43 | 0.718 | — |
| Breathing | Prefrontal | Low beta | −0.24 | 0.985 | — |
| Breathing | Prefrontal | High beta | −0.09 | 1.000 | — |

*Note.* $d_z$ = within-subject standardized effect size; positive = power increase, negative = decrease. $p_{FWER}$ = family-wise error rate from max-statistic sign-flip permutation across the complete 27-cell fast-band family (temporal, parietal, prefrontal × low beta, high beta, gamma × performance, mirroring, breathing vs. baseline). Cells with $p_{FWER} < 0.05$ ("Survived FWER" = Yes) are the 13 cells reproduced in Table 4. N = 12 per cell.

**Supplementary Table S5. Accelerometry Sensitivity Analysis for Intra-Brain High-Frequency Connectivity (Performance vs. Baseline)**

| Measure | HF change $d_z$ (Wilcoxon $p$) | Vigor association ρ ($p$) | Adjusted vigor slope $p$ | Movement-independent? |
|---|---|---|---|---|
| MIM (phase, leakage-robust) | 0.62 (.034) | +0.43 (.167) | .376 | Yes |
| AEC (orthogonalized amplitude) | 1.85 (.001) | +0.04 (.897) | .412 | Yes |
| aCOH (no leakage correction) | 0.66 (.052) | +0.62 (.031) | .017 | No (positive control) |

*Note.* HF = high frequency (mean of high-beta and gamma bands); vigor = change in mean dynamic-acceleration magnitude (performance − baseline). Effect sizes are Cohen's $d_z$ for the performance-versus-baseline change, with Wilcoxon signed-rank $p$. Associations with vigor are Spearman ρ; the adjusted slope $p$ is the unique contribution of vigor in a linear model of the connectivity change. Motion-pattern entropy was unrelated to MIM and AEC change ($\rho = -.41$ and $-.38$, both $p > .18$). AEC was elevated during mirroring ($d_z = 1.47$), where vigor did not rise, and was uncorrelated with vigor in that condition ($\rho = .32$, $p = .319$). aCOH's vigor association is consistent with its documented susceptibility to zero-lag source leakage and confirms the analysis detects movement when present. N = 12.

**Supplementary Table S6. Inter-brain coupling above the pseudo-dyad surrogate: full grid of all conditions, frequency bands, and connectivity measures.**

| Condition | Band | PLV *d* (*p*) | Coherence *d* (*p*) | Envelope *d* (*p*) |
|---|---|---|---|---|
| **Charlie Brown** | Theta (4–8) | −0.16 (.638) | 0.27 (.452) | **1.13 (.002)** |
| | Alpha (8–12) | −0.10 (.778) | 0.31 (.386) | **1.31 (.001)** |
| | Low beta (12–20) | −0.29 (.412) | 0.12 (.733) | **1.33 (.001)** |
| | High beta (20–30) | 0.01 (.978) | 0.56 (.110) | **1.29 (<.001)** |
| | Gamma (30–45) | −0.07 (.847) | **1.07 (.004)** | **1.39 (<.001)** |
| **Beethoven Day** | Theta (4–8) | −0.48 (.286) | −0.06 (.898) | **1.00 (.033)** |
| | Alpha (8–12) | −0.25 (.574) | −0.17 (.702) | **1.26 (.008)** |
| | Low beta (12–20) | −0.49 (.272) | −0.12 (.789) | **1.00 (.031)** |
| | High beta (20–30) | 0.03 (.949) | 0.39 (.385) | **1.11 (.015)** |
| | Gamma (30–45) | **0.91 (.046)** | **1.01 (.026)** | **0.95 (.037)** |
| **Happiness** | Theta (4–8) | −0.29 (.513) | −0.05 (.918) | 0.13 (.766) |
| | Alpha (8–12) | 0.87 (.056) | **0.90 (.048)** | 0.89 (.055) |
| | Low beta (12–20) | −0.48 (.287) | −0.39 (.380) | 0.13 (.768) |
| | High beta (20–30) | −0.72 (.109) | −0.59 (.194) | −0.67 (.135) |
| | Gamma (30–45) | 0.29 (.522) | 0.10 (.818) | **−1.01 (.029)** |
| **Mirror** | Theta (4–8) | 0.29 (.412) | 0.62 (.080) | **0.98 (.006)** |
| | Alpha (8–12) | 0.30 (.402) | 0.37 (.305) | 0.70 (.050) |
| | Low beta (12–20) | −0.01 (.978) | 0.14 (.683) | **1.57 (<.001)** |
| | High beta (20–30) | 0.06 (.874) | 0.56 (.118) | **1.59 (<.001)** |
| | Gamma (30–45) | −0.14 (.689) | **1.01 (.006)** | **1.71 (<.001)** |
| **Breathing** | Theta (4–8) | −0.36 (.303) | −0.54 (.130) | **−0.80 (.023)** |
| | Alpha (8–12) | −0.51 (.150) | −0.57 (.114) | −0.06 (.860) |
| | Low beta (12–20) | **−0.73 (.040)** | −0.49 (.161) | 0.44 (.215) |
| | High beta (20–30) | −0.01 (.968) | 0.11 (.761) | **0.84 (.018)** |
| | Gamma (30–45) | −0.41 (.246) | −0.29 (.411) | 0.53 (.130) |
| **Baseline** | Theta (4–8) | 0.01 (.973) | 0.04 (.908) | 0.00 (.994) |
| | Alpha (8–12) | −0.26 (.459) | −0.11 (.763) | 0.57 (.112) |
| | Low beta (12–20) | −0.40 (.263) | −0.49 (.162) | −0.23 (.520) |
| | High beta (20–30) | −0.22 (.524) | −0.30 (.398) | −0.19 (.582) |
| | Gamma (30–45) | 0.17 (.631) | −0.12 (.721) | −0.38 (.277) |
| **End-baseline** | Theta (4–8) | 0.21 (.632) | 0.53 (.239) | −0.02 (.971) |
| | Alpha (8–12) | −0.07 (.862) | 0.03 (.954) | −0.81 (.071) |
| | Low beta (12–20) | −0.11 (.815) | −0.23 (.630) | **−1.02 (.027)** |
| | High beta (20–30) | 0.07 (.871) | 0.11 (.797) | −0.08 (.859) |

| | | | | |
|---|---|---|---|---|
| | Gamma (30–45) | 0.17 (.700) | 0.03 (.947) | −0.02 (.958) |

*Note.* Values are Cohen's ***d*** (real vs. pseudo-dyad) with the two-sided permutation ***p*** (10,000 label-shuffle permutations) in parentheses. PLV = phase-locking value (phase only); coherence = combined phase and amplitude; envelope = amplitude-envelope correlation (amplitude only). Positive ***d*** indicates coupling exceeding the surrogate. Bold marks uncorrected ***p*** < .05; these ***p***-values are not corrected for multiple comparisons — the family-wise FDR-corrected ***q*** for the pre-specified primary family (three musical numbers × high beta and gamma × three measures) are reported in Table 4, Panel B. The pooled performance-average values appear in Table 4, Panel A. Real dyads ***n*** = 10 (Charlie Brown, mirror, breathing, baseline) or 7 (Beethoven Day, Happiness, end-baseline); pseudo-dyads ***n*** = 45 or 21, respectively (Table S3).

### Supplementary Table S7. Inter-brain amplitude coupling across all conditions

Real-versus-pseudo-dyad comparison (Cohen's *d*, pooled SD; permutation *p*, 10,000 label shuffles). Positive *d* indicates coupling in real co-performing dyads above same-condition surrogate dyads. Amplitude-envelope correlation and its leakage-robust orthogonalized variant, high-beta and gamma bands. $n = 10$ dyads.

| Condition | Env-corr high-beta | | Env-corr gamma | | Orth-AEC high-beta | | Orth-AEC gamma | |
|---|---|---|---|---|---|---|---|---|
| | *d* | *p* | *d* | *p* | *d* | *p* | *d* | *p* |
| Movement Mirror | +1.59 | <0.001 | +1.71 | <0.001 | +1.65 | <0.001 | +1.74 | <0.001 |
| Charlie Brown | +1.29 | <0.001 | +1.39 | <0.001 | +1.13 | 0.001 | +1.47 | <0.001 |
| Beethoven Day | +1.11 | 0.016 | +0.95 | 0.038 | +0.90 | 0.047 | +0.96 | 0.037 |
| Happiness (finale) | −0.67 | 0.135 | −1.01 | 0.029 | −0.52 | 0.250 | −1.13 | 0.015 |
| Paced Breathing | +0.84 | 0.019 | +0.53 | 0.132 | +0.80 | 0.025 | +0.58 | 0.107 |
| Rest baseline | −0.19 | 0.586 | −0.38 | 0.284 | −0.39 | 0.264 | −0.47 | 0.189 |
| End baseline | −0.08 | 0.856 | −0.02 | 0.960 | +0.04 | 0.925 | +0.02 | 0.970 |

Note. The graded pattern — movement mirroring, then the dyadic performance numbers (Charlie Brown, Beethoven Day), above ensemble and individual conditions (paced breathing, the Happiness finale), above rest — tracks the degree of interpersonal movement coordination each condition demands. Coherence, PLV, and imaginary coherence show the same ordering (not shown).

**Supplementary Table S8**

**Per-number decomposition of the performance effect across analysis layers (focal fast bands).**

| Layer / measure (domain) | Charlie Brown | Beethoven Day | Happiness | Pattern |
|---|---|---|---|---|
| PSD high-beta power (local) | +1.52 | +1.53 | +0.94 | Broad |
| PSD gamma power (local) | +1.50 | +1.70 | +0.94 | Broad |
| Intra-brain AEC – amplitude (within) | +1.96 | +2.61 | +1.41 | Broad |
| Intra-brain MIM – phase (within) | −0.01 | +1.92 | +0.29 | Beethoven-carried |
| Inter-brain envelope – amplitude (between) | +1.29 / +1.39 | +1.11 / +0.95 | −0.67 / −1.01 | Interaction-graded |
| Inter-brain orthog. AEC – amplitude (between) | +1.13 / +1.47 | +0.90 / +0.96 | −0.52 / −1.13 | Interaction-graded |

*Note.* Entries are effect sizes for the performance effect in the high-beta and gamma bands. For the local (PSD) and intra-brain rows, values are within-subject standardized change relative to baseline (dz; single value = mean of high beta and gamma). For the inter-brain rows, values are Cohen's d for real versus pseudo-dyads, shown as high beta / gamma. n varies by layer (PSD and intra-brain: 9–12 participants; inter-brain: 7–10 dyads). The pre-registered composite ("performance average," mean of the available musical numbers) reproduces the primary analyses: intra-brain MIM omnibus dz = 0.64, intra-brain AEC dz = 1.85. The table shows that local power and intra-brain amplitude coupling rise across all three numbers, whereas interaction-specific inter-brain coupling was selective — largest among the three numbers for Charlie Brown and, across all conditions, strongest during movement mirroring (Supplementary Table S10); intra-brain phase coupling (MIM), by contrast, is carried chiefly by Beethoven Day, underscoring that per-number attributions are measure-specific and provisional at this sample size.

**Supplementary Table S9. Spatial distribution of the amplitude-coupling effect across the 32 × 32 cross-brain channel grid, by condition and focal band.**

| Condition | Band | % pairs > surrogate | Mean *d* | Homol. *d* | Heter. *d* | MWU *p* | Top-10% share |
|---|---|---|---|---|---|---|---|
| **Charlie Brown** | High beta | 92% | 0.48 | 0.49 | 0.48 | .956 | 22% |
| | Gamma | 93% | 0.59 | 0.65 | 0.58 | .362 | 21% |
| **Beethoven Day** | High beta | 86% | 0.48 | 0.54 | 0.48 | .386 | 26% |
| | Gamma | 88% | 0.51 | 0.54 | 0.51 | .656 | 23% |
| **Happiness** | High beta | 37% | −0.14 | −0.16 | −0.14 | .681 | 59% |
| | Gamma | 24% | −0.30 | −0.27 | −0.30 | .880 | 78% |
| **Mirror** | High beta | 95% | 0.63 | 0.65 | 0.63 | .757 | 20% |
| | Gamma | 99% | 0.86 | 0.84 | 0.86 | .724 | 17% |
| **Breathing** | High beta | 71% | 0.19 | 0.21 | 0.19 | .703 | 32% |
| | Gamma | 68% | 0.16 | 0.10 | 0.16 | .385 | 33% |
| **Baseline** | High beta | 45% | −0.05 | 0.02 | −0.06 | .284 | 50% |
| | Gamma | 34% | −0.13 | −0.11 | −0.13 | .555 | 59% |
| **End-baseline** | High beta | 48% | −0.03 | −0.01 | −0.03 | .994 | 48% |
| | Gamma | 50% | −0.01 | −0.04 | −0.01 | .542 | 44% |

*Note.* Per-pair effects were computed as Cohen's ***d*** (real vs. pseudo-dyad) for each of the 1,024 cross-brain channel pairs. "% pairs > surrogate" is the proportion with ***d*** > 0. "Homol." (homologous) pairs are the 32 same-electrode cross-brain pairings (e.g., A-Cz with B-Cz); "Heter." (heterologous) are the remaining 992. MWU ***p*** is the Mann–Whitney comparison of homologous vs. heterologous ***d***. "Top-10% share" is the proportion of the total positive effect carried by the strongest 10% of pairs (an even spread across that decile would give ≈50%). The effect is broad and diffuse: real coupling exceeds surrogate at most pairs during the performance numbers and the mirror exercise, with no homologous-pair preference and no concentration in a small subset, whereas Happiness and the resting baselines fall at or below surrogate.

**Supplementary Table S10. Dyad roster and pseudo-dyad construction.**

| Dyad | Members (SVAEB_) | Member ages (yr) | Dyad mean age (yr) | Cohort | In 7-dyad conditions |
|---|---|---|---|---|---|
| 1 | 202 + 207 | 14, 9 | 11.5 | D1·G1 | **No (dropped)** |
| 2 | 202 + 213 | 14, 20 | 17.0 | D1·G1 | **No (dropped)** |
| 3 | 207 + 213 | 9, 20 | 14.5 | D1·G1 | **No (dropped)** |
| 4 | 203 + 209 | 14, 30 | 22.0 | D1·G2 | Yes |
| 5 | 203 + 210 | 14, 13 | 13.5 | D1·G2 | Yes |
| 6 | 209 + 210 | 30, 13 | 21.5 | D1·G2 | Yes |
| 7 | 205 + 206 | 19, 13 | 16.0 | D2·G1 | Yes |
| 8 | 201 + 208 | 24, 17 | 20.5 | D2·G2 | Yes |
| 9 | 201 + 212 | 24, 29 | 26.5 | D2·G2 | Yes |
| 10 | 208 + 212 | 17, 29 | 23.0 | D2·G2 | Yes |

*Note.* Each dyad comprises two participants who performed together. Cohort labels (D1/D2 × G1/G2) denote the production day and rehearsal group. Conditions performed by all participants (Charlie Brown, mirror, breathing, baseline) yield 10 dyads; conditions performed by only the second-day cohort (Beethoven Day, Happiness, end-baseline) yield 7 dyads, dropping the first three (cohort D1·G1). Pseudo-dyads were formed by pairing non-interacting participants within the same condition, giving 45 surrogate pairs for the 10-dyad conditions and 21 for the 7-dyad conditions. Because individuals recur across multiple pseudo-pairings, the surrogate samples are not mutually independent, and permutation ***p***-values are treated as approximate.
