## Supplementary Figures for "The Choreography of Coupled Brains: Coordinated Movement and Multi-Scale Neural Synchrony in Musical Theater for Individuals with Developmental Disabilities"

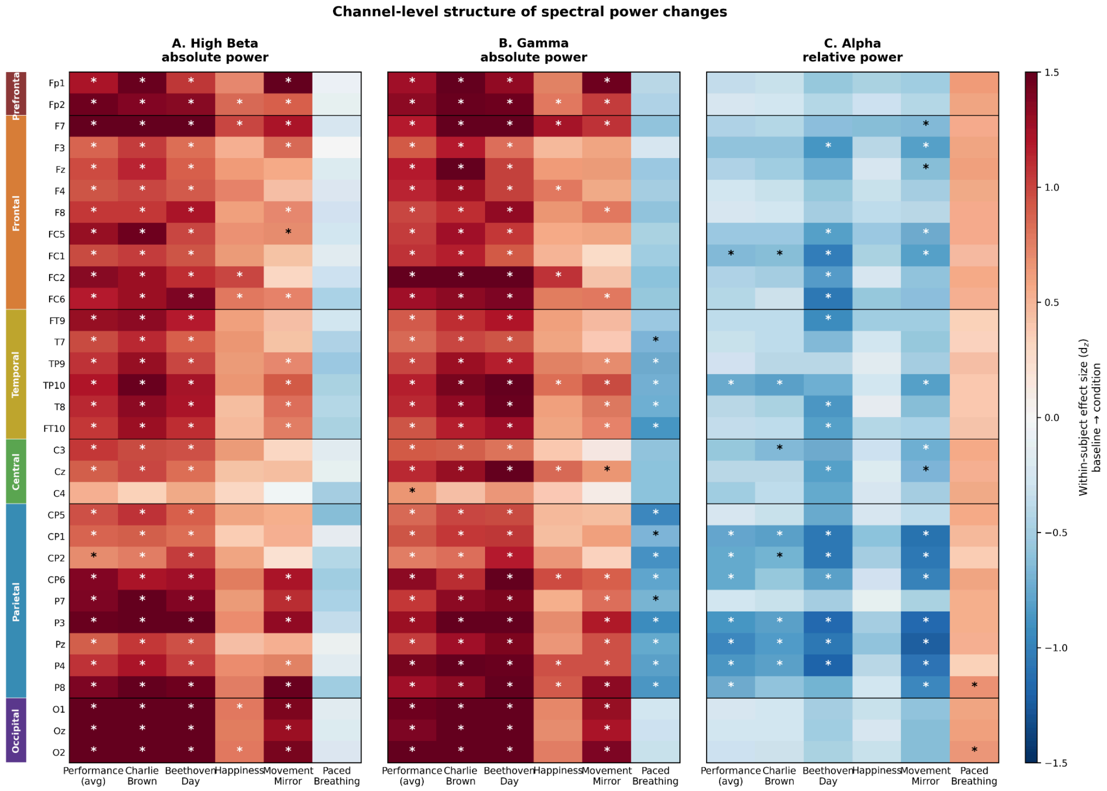


**Supplementary Figure S1.** *Channel-level structure of spectral power changes.* Per-channel within-subject effect sizes (d_z) for power change relative to resting baseline, shown for all 32 LiveAmp channels (rows, grouped by ROI on the left color bar) across six conditions (columns). Asterisks mark uncorrected $p < .05$ for each channel × condition cell. Panels A and B (high-beta and gamma absolute power) show the two fast-band anchors of Table 4. Active theatrical and movement conditions produce broad bilateral increases across nearly all channels; paced breathing shows the inverse pattern most prominently in gamma, with suppression spanning posterior–central electrodes (T7, TP10, CP1, P3, Pz, P4, P8). Panel C (alpha relative power) shows alpha as relative power (proportion of total delta–gamma power), the standard metric for the alpha-suppression / cortical-engagement literature. Absolute alpha is not displayed because the engagement construct refers specifically to redistribution of cortical activity away from the alpha rhythm: when fast-band power increases substantially in absolute terms, alpha proportion falls in relative terms even without an absolute alpha change. The relative metric isolates the proportional reallocation that constitutes the engagement signature, whereas absolute alpha conflates true alpha change with broadband scaling. Active conditions produce relative alpha suppression concentrated at parietal and centroparietal channels (CP1, CP2, P3, Pz, P4); paced breathing produces alpha increase strongest at O2 and P8. Statistical inferences remain at the ROI level (Table 2); channel-level effects are presented for descriptive context. N = 9 (Beethoven Day, mirror, breathing) or N = 12 (Charlie Brown, performance average).

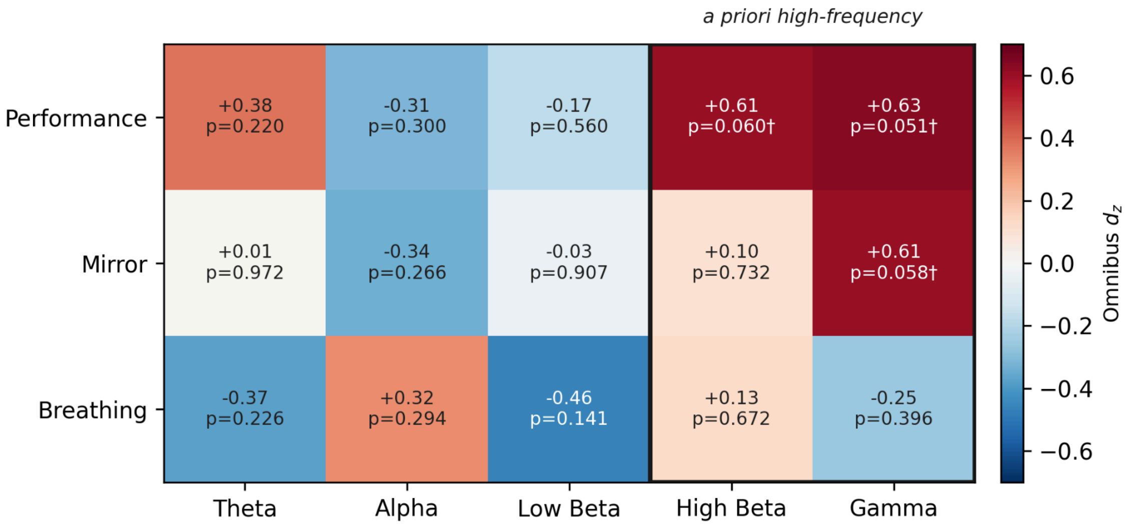


**Supplementary Figure S2.** *Network-omnibus effect size by condition and band.* Within-subject standardized effect size (d_z) for the per-participant network connectivity change (mean across all 28 Yeo network pairs) relative to resting baseline, for each condition (rows) and frequency band (columns); warm colors indicate increased coupling. The high-beta and gamma bands (boxed) constituted the a-priori high-frequency family. The performance-related increase was specific to the high-frequency bands and absent in theta, alpha, and low beta; movement mirroring showed an isolated gamma increase, and paced breathing showed no high-frequency increase. Cell annotations give d_z and the uncorrected one-sample t-test p-value against baseline († $p < .10$, * $p < .05$); these per-cell tests are descriptive, as the single confirmatory test was the combined high-frequency omnibus reported in the main text. n = 12.

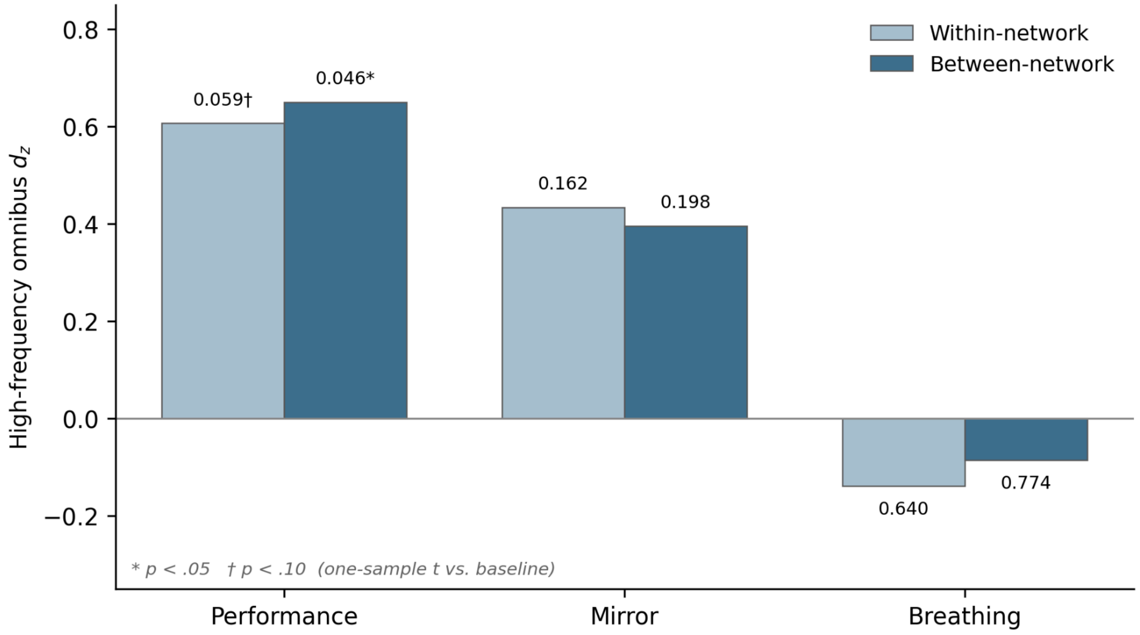


**Supplementary Figure S3.** *Within- versus between-network high-frequency change.* High-frequency (high-beta + gamma) omnibus effect size (d_z) for the per-participant connectivity change relative to baseline, decomposed into within-network (7 diagonal network pairs) and between-network (21 off-diagonal network pairs) components, by condition. The performance-related increase was present in both components and marginally larger for between-network coupling (d_z = 0.65, p = .046) than within-network coupling (d_z = 0.61, p = .059), consistent with increased cross-network integration. Annotations give the uncorrected one-sample t-test p-value against baseline (* p < .05, † p < .10). n = 12.

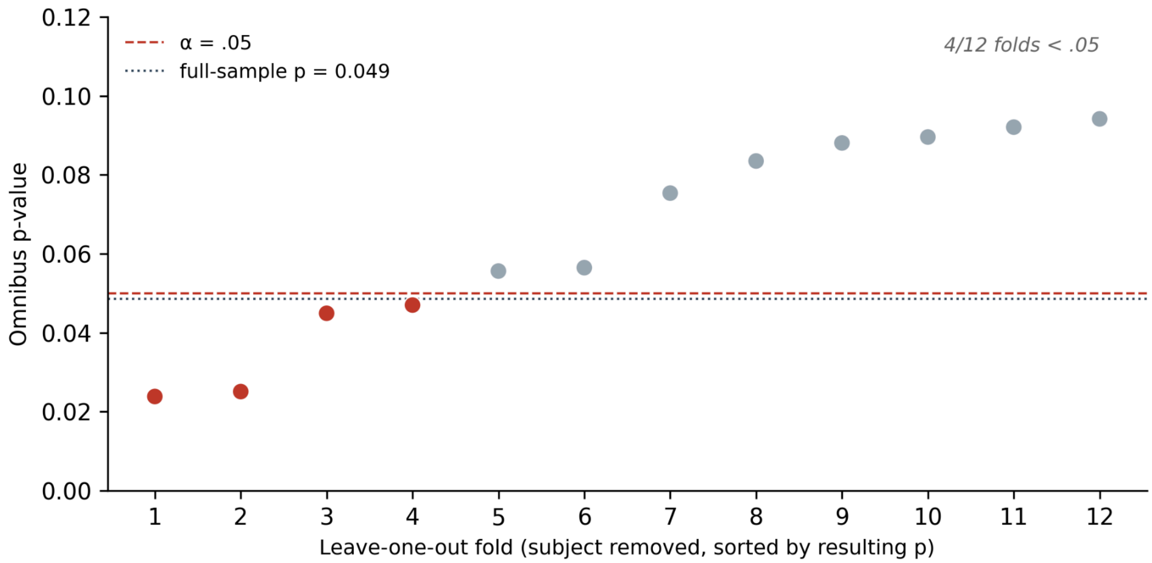


**Supplementary Figure S4.** *Leave-one-subject-out robustness of the performance high-frequency omnibus.* Omnibus p-value for the high-frequency performance effect when each participant is removed in turn, sorted by the resulting p-value. The full-sample result ($p = .049$; dotted line) falls just below $\alpha = .05$ (dashed line); under leave-one-out resampling the p-value ranged from .024 to .094 and remained below .05 in 4 of 12 folds. Red points indicate folds with $p < .05$, gray points folds with $p \geq .05$. The effect is present in the full sample but sensitive to individual participants, consistent with a small pilot. $n = 12$ (11 per fold).

A Orthogonalized cross-brain AEC (leakage-robust) · real vs. pseudo-dyad; links d > 1.0

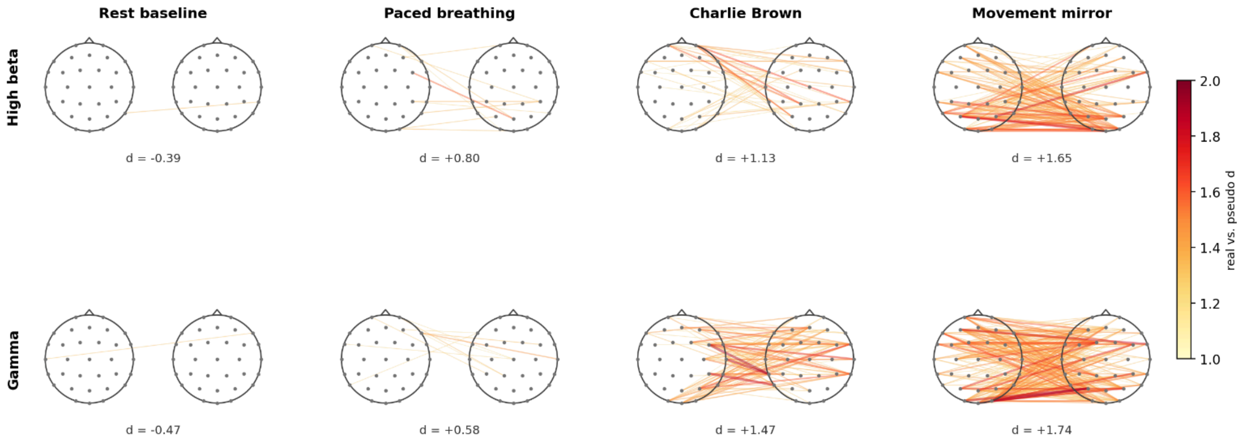


B Raw amplitude-envelope correlation (same gradient without orthogonalization) · links d > 1.0

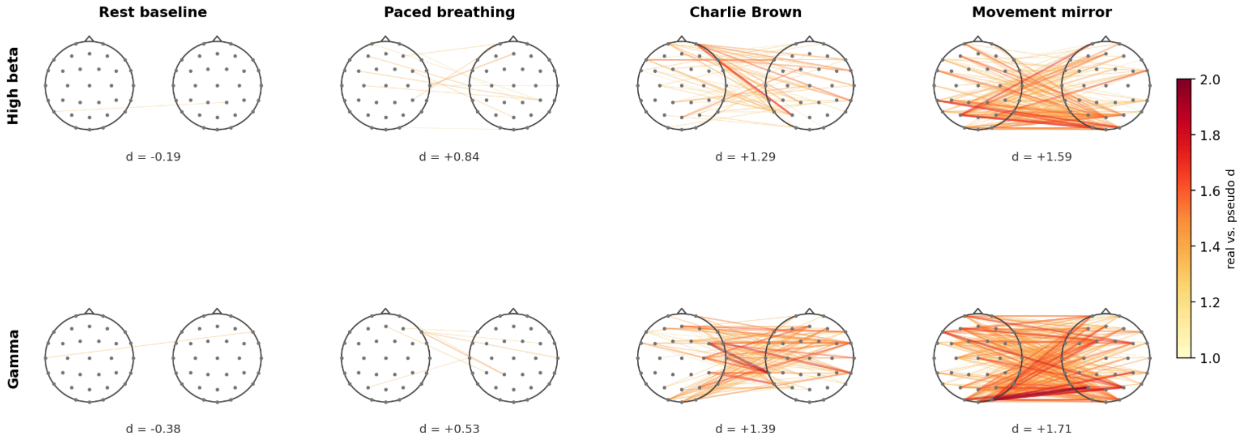


C Within-dyad increase in coupling from rest (orthogonalized AEC) · paired change vs. baseline; links d > 1.2

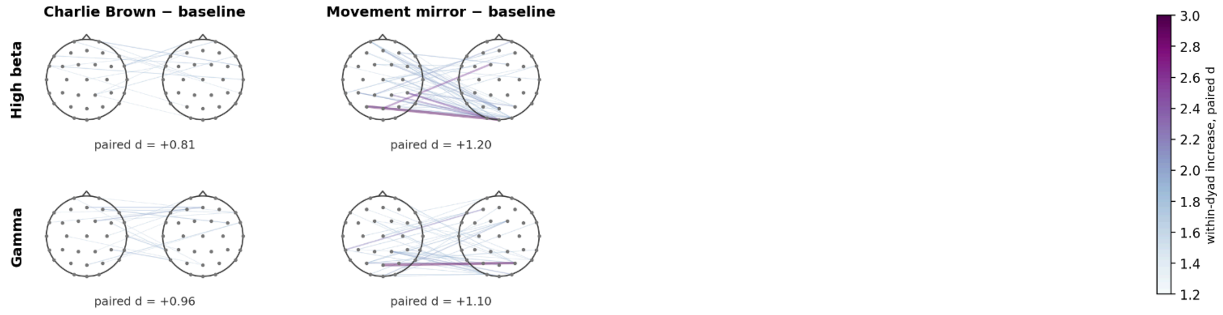

**Supplementary Figure S5. Inter-brain amplitude coupling scales with interpersonal movement coordination, with and without leakage correction.** Two-head renderings of cross-brain connectivity across the ten co-performing dyads. In each head pair, every line joins an electrode on one partner's scalp (Brain A, left) to an electrode on the other's (Brain B, right), with line color and width scaled by effect size; electrodes are shown in approximate 10–20 positions. **(A)** Absolute coupling by condition, computed with the orthogonalized cross-brain amplitude-envelope correlation (the leakage-robust amplitude measure) and expressed as the real-versus-pseudo-dyad standardized difference (Cohen's $d$). Lines show electrode pairs exceeding $d = 1.0$ on a color scale shared across all condition panels; the whole-brain $d$ is printed beneath each head pair. Coupling is near zero at rest and below the pseudo-dyad surrogate ($d = -0.39/-0.47$ for high-beta/gamma), rises during paced breathing (+0.80/+0.58), and is largest during the Charlie Brown performance (+1.13/+1.47) and, most strongly, the movement-mirroring exercise (+1.65/+1.74). **(B)** The identical analysis computed with the raw amplitude-envelope correlation (without orthogonalization), on the same threshold and color scale. The coordination gradient is essentially unchanged (whole-brain $d$: rest ≈ 0, breathing +0.84/+0.53, Charlie Brown +1.29/+1.39, mirror +1.59/+1.71), indicating that it does not depend on the leakage correction. **(C)** The within-dyad increase in coupling from rest, computed as the paired standardized change from baseline (Charlie Brown or mirror minus baseline across dyads, orthogonalized measure); lines show pairs exceeding a paired $d = 1.2$. Coupling grows within the same dyads during both active conditions, more so during mirroring (whole-brain paired $d = +0.81/+0.96$ for Charlie Brown, +1.20/+1.10 for mirror). This figure is provided for visualization; formal inference (per-condition effect sizes, pseudo-dyad permutation tests, and leave-one-out robustness) is reported for the focal bands in Figure 4 and Table 5. $n = 10$ dyads per condition; electrode positions reflect the recording montage.

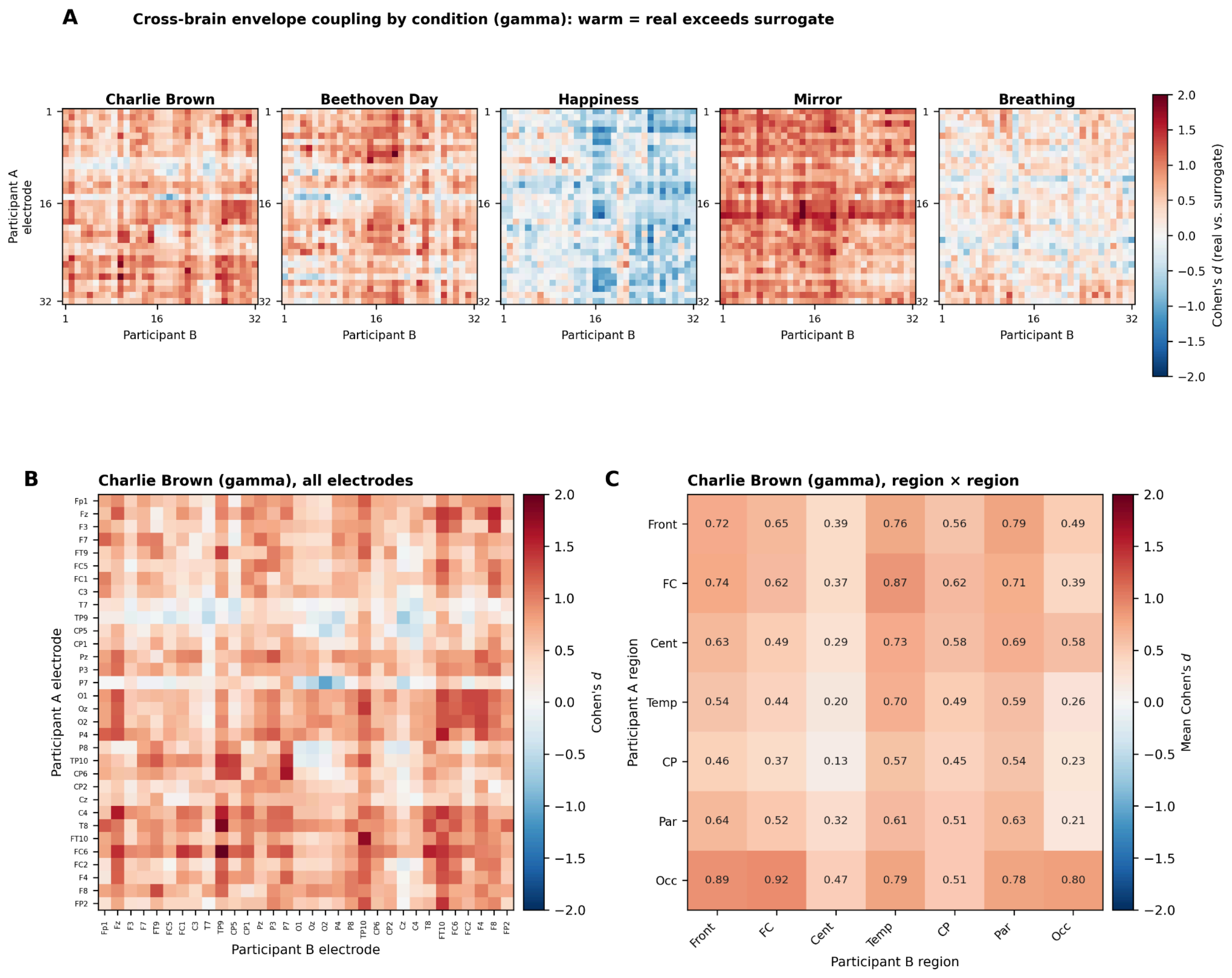


**Supplemental Figure S6.** *Spatial distribution of inter-brain amplitude-envelope coupling.* (A) Cross-brain coupling maps (Cohen's ***d***, real vs. pseudo-dyad) in the gamma band for each condition; rows index participant A's 32 electrodes, columns participant B's. Warm values indicate real coupling exceeding surrogate. The performance numbers (Charlie Brown, Beethoven Day) and the mirror exercise are broadly warm; Happiness is broadly cool (below surrogate), consistent with its shared-stimulus structure inflating the pseudo-dyad baseline; breathing is weak. (B) The Charlie Brown gamma map at full resolution with all 32 electrode labels. (C) The same data averaged within a seven-region anatomical parcellation (region × region mean ***d***). Coupling is positive across all 49 region pairs (***d*** = 0.13–0.92), with within-region coupling (mean ***d*** = 0.60) essentially equal to between-region coupling (mean ***d*** = 0.55), indicating a diffuse effect with no concentration in a specific network; the modestly higher values involving temporal and occipital sites cannot be distinguished from muscle contribution without the planned movement control. Front = frontal, FC = fronto-central, Cent = central, Temp = temporal, CP = centro-parietal, Par = parietal, Occ = occipital.

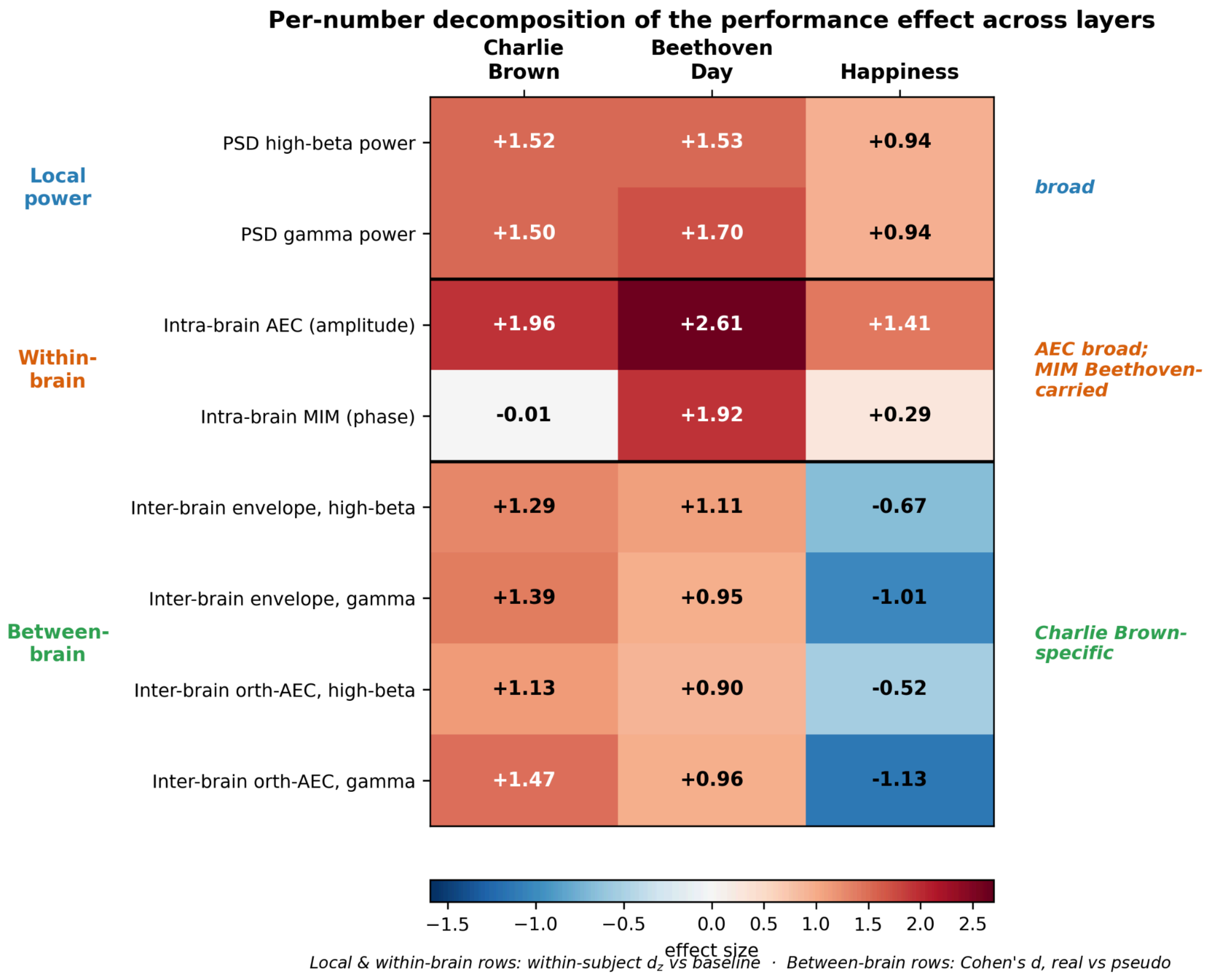


**Supplementary Figure S7.** *Per-number decomposition of the performance effect across analysis layers.* Effect sizes for the change during each of the three musical numbers (columns: Charlie Brown, Beethoven Day, Happiness) for eight measures grouped by analysis layer (rows). Cell color and the annotated value give the effect size (diverging scale centered at zero; warm = power increase or coupling above surrogate, cool = decrease or coupling below surrogate). The metric differs by layer and is not directly comparable across the horizontal rules. For the two local-power rows (per-channel spectral power) and the two within-brain rows (source-space intra-brain connectivity), values are within-subject standardized change relative to resting baseline (*d*_z; *n* = 9–12 participants). For the four between-brain rows (inter-brain connectivity), values are Cohen's *d* for real versus same-condition pseudo-dyads (*n* = 7–10 real dyads and 21–45 pseudo-dyads). The two local-power rows are shown separately for high beta and gamma; the within-brain rows summarize the high-frequency band (high beta and gamma) for the orthogonalized amplitude-envelope measure (AEC) and the phase measure (MIM); the four between-brain rows show the raw amplitude-envelope correlation and the leakage-robust orthogonalized cross-brain AEC, each separately for high beta and gamma. The pre-registered composite ("performance average," the mean of the available numbers) reproduces the primary analyses reported in the main text (intra-brain MIM omnibus *d*_z = 0.64; intra-brain AEC *d*_z = 1.85). The figure illustrates a layer-dependent dissociation: local fast-band power and within-brain amplitude coupling increase across all three numbers (broad); the within-brain phase measure (MIM) is carried chiefly by Beethoven Day, with Charlie Brown near zero; and interaction-specific between-brain coupling is confined to Charlie Brown — present for Beethoven Day and absent (below surrogate) for the full-ensemble Happiness finale. AEC = amplitude-envelope correlation; MIM = multivariate interaction measure; PSD = power spectral density; *d*_z = within-subject standardized effect size.
