## Supplementary Methods and Results for "The Choreography of Coupled Brains: Coordinated Movement and Multi-Scale Neural Synchrony in Musical Theater for Individuals with Developmental Disabilities"

### *Details and Psychometric Properties of Neuropsychological Assessments*

Revised Children's Anxiety and Depression Scale (RCADS)(Chorpita *et al.*, 2000). The RCADS is a 47-item self-report questionnaire assessing six DSM-aligned anxiety and depression dimensions: Separation Anxiety, Social Phobia, Generalized Anxiety, Panic Disorder, Obsessive-Compulsive Disorder, and Major Depression. Respondents rate each item as "never" (0), "sometimes" (1), "often" (2), or "always" (3). The first five subscales sum to a Total Anxiety scale, and all six to a Total Internalizing scale; we designated Total Anxiety as the primary behavioral outcome. In the original youth validation, the RCADS demonstrated excellent internal consistency (Total Internalizing $\alpha$ = 0.93–0.96) and satisfactory test–retest reliability and concurrent validity(Chorpita *et al.*, 2000).

Piers-Harris Self-Concept Scale, Third Edition (Piers-Harris 3)(Piers *et al.*, 2002). The Piers-Harris 3 is a 58-item self-report measure of global and domain-specific self-concept, normed for ages 6–22 and written at approximately a first-grade reading level. Items are scored 0 (negative self-concept) or 1 (positive self-concept) and yield a Total Score and six domain scores: Behavioral Adjustment, Intellectual and School Status, Physical Appearance and Attributes, Freedom from Anxiety, Popularity, and Happiness and Satisfaction. Internal consistency in the standardization sample was high for the Total Score ($\alpha \approx 0.91$) and acceptable to good for the six domain scales ($\alpha$ = 0.74–0.83).

KIDSCREEN-52 Quality of Life Questionnaire (KIDSCREEN-52)(Ravens-Sieberer *et al.*, 2008). Health-related quality of life was assessed using the KIDSCREEN-52, a 52-item self-report questionnaire covering ten domains: Physical Well-Being, Psychological Well-Being, Moods and Emotions, Self-Perception, Autonomy, Parent Relations and Home Life, Social

Support and Peers, School environment, Social Acceptance (Bullying), and Financial Resources. Items are rated on a five-point Likert scale from 1 ("never"/"not at all") to 5 ("always"/"extremely"), with negatively worded items reverse-coded, and higher scores indicating better health-related quality of life. The KIDSCREEN-52 has demonstrated good internal consistency (α = 0.77–0.89) and satisfactory test–retest reliability (ICCs = 0.56–0.77).

Social Connectedness Scale–Revised (SCS-R)(Lee *et al.*, 2001). The SCS-R is a 20-item self-report questionnaire assessing perceived interpersonal connectedness. Items are rated on a six-point Likert scale from 1 ("strongly disagree") to 6 ("strongly agree"); items are phrased both positively and negatively to minimize response bias, with negatively worded items reverse-scored. The SCS-R has demonstrated excellent internal consistency (α = 0.92) and strong two-week test–retest reliability ($r$ = 0.94).

Emotional Regulation Strategies for Artistic Creative Activities Scale (ERS-ACA)(Fancourt *et al.*, 2019). The ERS-ACA is an 18-item self-report measure of emotion regulation strategies used during artistic creative activities, with items grouped into three subscales - Avoidance, Approach, and Self-Development - rated on a five-point Likert scale from 1 ("strongly disagree") to 5 ("strongly agree"). Internal consistency is strong (general factor α = 0.93; subscales α = 0.88–0.90), with good test–retest reliability for the general factor and the approach and self-development subscales (ICCs = 0.75–0.90) and moderate reliability for avoidance (ICC = 0.50–0.75). The ERS-ACA was administered during the pre-test as a baseline predictor of intervention response.

Qualitative Open-Ended Assessments: At pre-test, participants answered two open-ended questions: (1) "How would you describe your recent thoughts and feelings? By recent, we mean within the past three months," and (2) "Describe your relationship with the STEP VA community." At post-test, they answered three: (1) "How would you describe your recent thoughts and feelings? By recent, we mean during the past month, particularly focusing on the time you have been rehearsing with the STEP VA community," (2) "Describe your relationship with the STEP VA community. Has your relationship changed since you started rehearsing for the musical?" and (3) "Is there anything else you would like to share about your experience rehearsing for or performing in the *Charlie Brown* musical?" Parents/guardians answered the same questions from their perspective of their child's experiences. These items were investigator-developed and not formally validated.

***Qualitative Analysis***

Open-ended responses were analyzed using reflexive thematic analysis following the six-phase framework of Braun and Clarke (2006), selected for its flexibility, compatibility with small qualitative datasets, and suitability for applied community-based research(Braun and Clarke, 2006). All four datasets (participant and parent/guardian, pre and post) were read in full across multiple familiarization passes, after which meaningful units were coded inductively, attending to both explicit content and latent meaning, and clustered into candidate themes that were iteratively reviewed, refined, and named using participants' own language where possible. The primary analytic frame was group-level comparison: pre- and post-responses were examined across the full sample of each respondent group to document patterns in emotional tone, response specificity, and thematic content, accommodating all respondents regardless of whether they completed surveys at both time points. Where respondents provided data at both time points (n = 10 participants; n = 9 parents/guardians), individual-level matched-pair comparison was conducted as a secondary layer to examine whether group-level patterns were consistent with individual trajectories and to contextualize outlier responses. Convergence of themes across respondent groups was treated as a strengthening indicator of salience, and an audit trail of codes and analytic memos was maintained throughout to support rigor(Lincoln and Guba, 1988).

#### ***Details of EEG Analysis***

*EEG Acquisition and Preprocessing*

Mobile EEG was recorded with 32-channel wet-electrode caps (LiveAmp 32, Brain Products GmbH, Gilching, Germany); participants were recorded in groups of one to three. During the breathing exercise, participants inhaled for four counts, held for two, and exhaled for four, following the instructor's lead; during the mirroring task, they replicated the instructor's movements, a technique used in prior rehearsals to prepare for performance. Data from all conditions were appended in sequential order for each participant before preprocessing in EEGLAB(Delorme and Makeig, 2004). Recordings were bandpass-filtered between 1 and 55 Hz using a zero-phase Hamming-windowed FIR filter. Channels that were flat for more than 5 s, exceeded 4 standard deviations of the remaining channels in noise, or correlated below 0.8 with neighboring channels were removed automatically (an average of 2.9 channels were interpolated per dataset; SD = 1.8, max = 6). Portions of data varying more than 20 standard deviations from artifact-free segments were then corrected by Artifact Subspace Reconstruction through a 0.5-s PCA(Chang *et al.*, 2018). Channels removed before ASR were interpolated, and the data were full-rank re-referenced to the common average. Datasets were decomposed by Adaptive Mixture Independent Component Analysis (AMICA; single model, 2000 iterations)(Hsu *et al.*, 2018), which iteratively rejects artifact-contaminated segments during fitting (rejection criterion 3 SD; one outlier removed for every 15 iterations). Independent components were then classified by

ICLabel(Pion-Tonachini *et al.*, 2019). Two component-inclusion criteria were applied: for power spectral density analysis, only components labeled over 50% brain were retained(Gonsisko *et al.*, 2023); for intra- and inter-brain connectivity analyses, components labeled over 90% muscle, eye, heart, line-noise, or other artifact were removed. After pruning, each participant's dataset was split into separate datasets per condition while preserving the estimated component weights. Because source reconstruction and multivariate connectivity estimation can be destabilized when data are strongly rank-deficient, each artifact-corrected recording was screened for rank sufficiency before analysis. Following component rejection and re-referencing, the effective rank of every 32-channel recording was computed as the number of singular values exceeding a relative tolerance of $1 \times 10^{-6}$ (normalized to the largest singular value). Because some reduction below the 32 recorded channels is expected after average referencing, interpolation, and component removal, recordings were retained only if their effective rank met a pre-specified floor of 15 and contained no flat or non-finite channels. To confirm that inclusion did not depend on this threshold, the screen was repeated at floors of 12 and 18.

*Computing Power Spectral Density (PSD)*

Power spectral density was computed from channel-level time series. Each continuous recording was segmented into fixed-length epochs with 50% overlap; epoch length was chosen by condition to balance frequency resolution against signal stationarity: four-second windows for seated, movement-free conditions (resting baseline, end-of-session baseline, and paced breathing), two-second windows for standing movement mirroring, and one-second windows for the theatrical performance conditions. For each channel and epoch, the signal was tapered with a Hann window and transformed by Fast Fourier Transform (length $2\char`^\lceil \log_2 N \rceil$). One-sided power spectra were scaled (doubling non-DC, non-Nyquist bins), normalized by the window's summed squared coefficients, and averaged across epochs in Welch fashion, which preserves the non-phase-locked oscillatory activity that dominates resting EEG.

Mean spectral power was then computed within six non-overlapping frequency bands following a prespecified band scheme - delta (1–4 Hz), theta (4–8 Hz), alpha (8–12 Hz), low beta (12–20 Hz), high beta (20–30 Hz), and gamma (30–45 Hz) - and additionally within a broadband beta (12–30 Hz) used only for band-ratio computations and excluded from total-power normalization to avoid double-counting low and high beta. Power was expressed in two metrics: absolute power, the $\log_{10}$-transformed band-averaged power (log $\mu V^2/Hz$), and relative power, the proportion of total power across the six non-overlapping bands. Confirmatory analyses within the pre-registered fast-band family used absolute power, which directly indexes oscillatory activity in each band and is not subject to the closed-system constraint of relative metrics. Exploratory analyses of alpha suppression used relative power, because the alpha-suppression construct as defined in the cortical-engagement literature(Mathewson *et al.*, 2012) mechanistically refers to a proportional redistribution of activity away from the alpha rhythm and is conventionally indexed as a proportion of total power. We do not report absolute alpha because broadband increases during active conditions can mask the proportional reallocation that the engagement construct concerns; the relative metric isolates this reallocation directly. Both metrics appear in the supplemental tables for transparency.

Region-of-interest (ROI) power was computed by averaging channel power within six anatomical groupings defined by 10-20 channel labels: prefrontal (Fp1, Fp2), frontal (F3, F4, F7, F8, Fz, FC1, FC2, FC5, FC6), central (C3, C4, Cz), temporal (T7, T8, FT9, FT10, TP9, TP10), parietal (P3, P4, P7, P8, Pz, CP1, CP2, CP5, CP6), and occipital (O1, O2, Oz). Channel-to-ROI assignment was made by label, so ROI values reflect averages of the named channels regardless of their row order in the underlying data matrix. Based on a priori hypotheses about the engagement of frontal, posterior, and association cortex during musical theater performance, the confirmatory inferential analyses focused on the temporal, parietal, and prefrontal ROIs in the three fast bands (low beta, high beta, gamma), constituting a pre-registered family of 27 cells (3 ROIs × 3 bands × 3 conditions), in which each program condition - performance, movement mirroring, and paced breathing - was contrasted against resting baseline. All six ROIs and seven bands appear in the descriptive supplemental tables. Conditions included resting baseline (reference), end-of-session baseline, paced breathing, movement mirroring, and the four theatrical performance conditions (*Charlie Brown*, *Beethoven Day*, *Happiness*, and a per-subject performance-average composite across these).

*Computing Functional Connectivity (Intra-brain synchrony)*

Source-level functional connectivity was computed using a preprocessing pipeline parallel to that used for spectral power, with two adjustments suited to connectivity estimation. First, because connectivity is degraded by overly aggressive component removal(Castellanos and Makarov, 2006), a conservative ICLabel criterion(Pion-Tonachini *et al.*, 2019) was applied: independent components were removed only when classified as at least 90% likely to reflect muscle, eye, heart, line-noise, or "other" (non-brain) activity, retaining as many brain components as possible for source reconstruction. Second, to maximize the number of components available for the ICA decomposition during movement-rich conditions, all activities were concatenated into a single dataset per participant for preprocessing and decomposition, then separated by activity before connectivity estimation.

Equivalent current dipoles were fit to each retained independent component using EEGLAB's DIPFIT plugin against a boundary element head model derived from the Montreal Neurological Institute template brain. Electrode coordinates were co-registered to the head model using the following transformation: translation [0.94, −16.90, 1.46] mm, rotation [0, 0, −1.5708] radians, and unit scaling. Components whose scalp topography was bilaterally symmetric were re-fit as two symmetric dipoles using the fitTwoDipoles plugin(Piazza *et al.*, 2016).

For each activity dataset, a lead field was computed from the Colin27 template head model (head_modelColin27_5003_Standard-10-5-Cap339) included in EEGLAB. Sensor-level data were projected to source space using a linearly constrained minimum-variance (LCMV) beamformer and downsampled to 100 Hz, as implemented in the ROIconnect plugin(Pellegrini *et al.*, 2023), and parcellated into the 68 cortical regions of the Desikan-Killiany atlas(Desikan *et al.*, 2006). To separate the phase and amplitude contributions to intra-brain coupling, three connectivity measures were estimated from this common source reconstruction. The pre-registered primary measure was the multivariate interaction measure (MIM)(Ewald *et al.*, 2012; Marzetti *et al.*, 2013), a multivariate generalization of the imaginary part of coherency(Nolte *et al.*, 2004) that is insensitive to zero-lag (volume-conduction) artifacts; MIM was computed from the three dominant principal components per region via a multivariate autoregressive model (model order = 20). To isolate amplitude coupling, orthogonalized amplitude-envelope correlation (AEC) was computed from the single dominant source signal per region: signals were band-pass filtered, Hilbert-transformed, and pairwise-orthogonalized before their amplitude envelopes were correlated(Brookes *et al.*, 2012; Hipp *et al.*, 2012). Orthogonalization is required within a single brain, where amplitude-envelope correlation is otherwise inflated by zero-lag leakage. As a combined phase-and-amplitude reference, absolute coherence (aCOH) was computed from the same three-component source model; because aCOH retains the zero-lag component, it remains susceptible to the volume-conduction leakage that MIM and orthogonalized AEC remove, and it is therefore reported only alongside MIM rather than as a standalone result. Each measure was estimated across the spectrum and averaged within five frequency bands - theta (4–8 Hz), alpha (8–12 Hz), low beta (12–20 Hz), high beta (20–30 Hz), and gamma (30–45 Hz) - yielding a 68 × 68 region-by-region matrix per measure, participant, condition, and band. For each measure, a performance-average matrix was additionally computed per participant as the mean of the cb, bd, and happy performance matrices.

For the primary (MIM) analysis, to test whether musical-theater engagement alters intra-brain synchrony at the level of large-scale functional systems, each 68 × 68 region-by-region MIM matrix was aggregated to the seven canonical Yeo networks (visual, somatomotor, dorsal attention, ventral attention/salience, limbic, frontoparietal control, and default mode)(Yeo *et al.*, 2011). Each Desikan-Killiany region was assigned to its dominant Yeo network using a fixed, published spatial-overlap mapping; network-pair connectivity was computed as the mean MIM across all region pairs spanning the two networks, with within-network values excluding the zero diagonal. This produced a 7 × 7 network connectivity matrix (7 within-network and 21 between-network values) per participant, condition, and frequency band. The two secondary measures (orthogonalized AEC and aCOH) were instead summarized at the whole-brain level, as the mean across all region pairs, the scale at which all three measures are directly comparable. For each active condition (performance average, movement mirroring) and the paced breathing condition, connectivity change was computed as the signed difference from resting baseline, retaining participants with paired data.

*Computing inter-brain synchrony*

As an exploratory aim, we examined inter-brain synchrony during shared performance using hyperscanning connectivity computed in HyPyP(Ayrolles *et al.*, 2021), applied to the same preprocessed 32-channel data. Each performance group was recorded simultaneously on hardware-synchronized amplifiers (shared synchronization-pulse trains across amplifier pairs), so the two members' per-condition files derived from a common time base and their epochs were temporally aligned across partners. For each dyad (two participants who performed together in the same group) the two members' recordings for a given condition were segmented from recording onset into 4-s non-overlapping epochs; a fixed 4-s window was used across all conditions to provide sufficient cycles for stable estimation in the lower bands, and the two members' epoch series were truncated to a common length. Complex spectra were estimated within five frequency bands (theta, 4–8 Hz; alpha, 8–12 Hz; low beta, 12–20 Hz; high beta, 20–30 Hz; gamma, 30–45 Hz). The delta band (< 4 Hz) was not analyzed: the 4-s epochs provide too few low-frequency cycles for stable delta estimation, and delta is especially susceptible to movement and other non-neural artifact in mobile recordings; this five-band scheme also matches the spectral-power and intra-brain connectivity layers. Five connectivity measures were derived from the same complex signal and averaged across epochs. Three index coupling without separating instantaneous from lagged components: coherence (sensitive to both phase and amplitude consistency), the phase-locking value (PLV; phase only)(Lachaux *et al.*, 1999), and amplitude-envelope correlation (amplitude only). Two are zero-lag-insensitive analogues: imaginary coherence(Nolte *et al.*, 2004), which retains only the lagged phase relationship, and an orthogonalized cross-brain amplitude-envelope correlation, which pairwise-orthogonalizes the two participants' band-limited analytic signals to remove their instantaneously shared component before correlating amplitude envelopes(Brookes *et al.*, 2012; Hipp *et al.*, 2012). The interpretation of the zero-lag component differs between the two layers: within a single brain it reflects volume conduction, whereas between two brains it reflects instantaneously shared input - most importantly the common audiovisual stimulus and any correlated movement of the two partners. The orthogonalized measure therefore provides the critical test of whether inter-brain amplitude coupling reflects genuine lagged coupling rather than a shared driver. Deriving all five measures from one complex signal allowed the phase and amplitude contributions to inter-brain coupling, and their zero-lag-insensitive counterparts, to be separated. For each measure and band, the cross-brain block of the connectivity matrix (participant-1 channels × participant-2 channels; 32 × 32) was retained, and its mean indexed whole-montage inter-brain connectivity for that dyad.

Because two people responding to the same stimulus can show elevated inter-brain connectivity without any genuine interaction, a pseudo-dyad surrogate control was used to separate interaction-related coupling from coupling attributable to shared task structure. Pseudo-dyads were formed by pairing participants who completed the same condition but belonged to different performance groups, and who therefore never performed together; their connectivity was computed identically. Real-dyad connectivity exceeding the pseudo-dyad distribution indicates coupling beyond that produced by the shared stimulus. Usable data yielded 10 real dyads for the resting baseline and the *Charlie Brown*, mirror, and breathing conditions, and 7 for the *Beethoven Day, Happiness,* and end-baseline conditions (one group lacked usable segments for the latter three); the corresponding pseudo-dyad samples comprised 45 and 21 cross-group pairs. A performance average was computed for each dyad as the mean of the three musical numbers (*Charlie Brown, Beethoven Day, Happiness*), restricted to dyads with usable data for all three.

**Sensitivity Analyses**

*Age-covariate sensitivity.* Because participants spanned a wide age range (9–30 years), and because both spectral power and functional connectivity change substantially across development, all confirmatory and primary neural effects were re-examined with age as a covariate. For each affected measure — the spectral-power cells surviving family-wise correction, the high-frequency intra-brain connectivity omnibus, and the inter-brain coherence contrasts — the change score was regressed on mean-centered age by ordinary least squares; the intercept indexed the age-adjusted mean change at the sample mean age, and the slope tested whether the change varied with age, with the zero-order association additionally examined by Spearman correlation. Participant-level regressions used mean-centered participant age; the inter-brain measures, defined at the level of dyads, used mean-centered dyad mean age (the mean of the two members' ages). These age-adjusted analyses are reported alongside the unadjusted results as sensitivity checks throughout.

*Movement sensitivity.* Because the active conditions entailed overt movement, every spectral and connectivity effect surviving family-wise correction, together with the exploratory alpha-suppression effects, was additionally tested for dependence on movement. Movement was indexed by accelerometric vigor — the mean dynamic-acceleration magnitude of the head-mounted sensor within each segment — computed by per-axis high-pass filtering at 0.25 Hz to remove the gravitational component, combining the three axes into a single dynamic-acceleration magnitude, and decimating to 50 Hz. For each participant, vigor was averaged across the performance segments and across the resting baseline, and a performance-minus-baseline vigor change score was formed in parallel to the neural change scores; the inter-brain measures used the dyad-mean vigor change. Movement dependence was evaluated in two ways: (i) the per-participant (or per-dyad) change in the neural measure was correlated with the concurrent change in vigor using Spearman's ρ; and (ii) the effect was re-estimated with vigor entered as a linear covariate, with movement treated as a potential confound if the vigor slope was significant or if adjustment materially attenuated the effect.

*Spectral parameterization.* To determine whether any movement-related variance in the spectral-power effects resided in oscillatory or in broadband activity, the power spectra were decomposed into an aperiodic (1/f-like) component and a set of periodic peaks using spectral parameterization (specparam)(Donoghue *et al.*, 2020). Spectra were fit from 2 to 44 Hz in the fixed aperiodic mode (peak-width limits 1–8 Hz; maximum six peaks; minimum peak height 0.05), with the upper bound placed below the 45-Hz low-pass roll-off of the recordings. The aperiodic offset and exponent, and the summed periodic peak power within each canonical band, were extracted for every participant, condition, and channel, and the two movement tests above were repeated separately on the aperiodic and periodic components. The same parameterization was applied at the source level to the dipole-clustered independent components. Model fit was high at the source level ($R^2 \approx 0.96$–$0.99$) but poor for a minority of scalp channels (≈ 21% with $R^2 < 0.90$), with oscillatory peaks detectable in only a minority of channel spectra at this sample size; the parameterization is therefore reported as a sensitivity analysis rather than a primary outcome.

### SUPPLEMENTARY RESULTS

#### Behavioral: exploratory ER predictors

For participants, higher baseline general emotion-regulation, approach-strategy, and self-development scores were associated with larger quality-of-life gains (KIDSCREEN Total change: General Factor $\rho = 0.93$, $p = 0.008$; Approach $\rho = 0.90$, $p = 0.015$; Self-Development $\rho = 0.88$, $p = 0.021$; all $n = 6$). With $n = 6$ these are hypothesis-generating only and are not interpreted further. Full ERS-ACA correlation grid in Supplementary Table S2.

**Behavioral: rater agreement**

Baseline agreement between raters was strong for the two measures with the largest pre-/post-effects: RCADS Total Anxiety (Spearman $\rho = 0.80$, $n = 12$, $p = 0.002$) and Piers-Harris Total ($\rho = 0.78$, $n = 12$, $p = 0.003$), indicating that participants and parents began with convergent views of anxiety severity and self-concept. Agreement on change scores ($n = 7$) was weaker and, for self-concept, divergent: RCADS Total Anxiety change agreement was moderate ($\rho = 0.56$, n.s.), whereas Piers-Harris change agreement was slightly negative ($\rho = -0.21$). This combination — convergent baselines but divergent change — quantifies the rater discrepancy and is consistent with raters sharing a view of the child's starting state but responding to different cues when judging improvement. Given $n = 7$ for the change-score correlations and their instability under leave-one-out resampling, these agreement estimates are descriptive only (full statistics in Supplementary Table S3).

**Behavioral: qualitative themes**

The most prevalent theme across open-ended responses was belonging and identity (e.g., "his people," "my people," "feels like family"). A second theme was competence through challenge ("He was challenged and he did it — we are proud and he is proud of himself"). Among a small number of parents ($n = 2$), a theme was that participants were perceived differently by others through their performances. Further themes across participants and parents were that the program built a community-sustaining infrastructure for families and gave participants a sense of purpose ("This is the first thing that has given [him] a purpose").

**PSD: age covariate**

Because participants spanned a wide age range (9–30 years), each surviving cell was re-tested with age as a covariate (per-subject change regressed on mean-centered age). All 13 family-wise survivors remained significant after adjustment (all adjusted $p \leq 0.006$), no cell showed a significant association between power change and age (age-slope $p = 0.07$–$0.97$), and effect sizes were essentially unchanged. Age was therefore not a meaningful confound of the spectral-power findings.

**PSD: spectral parameterization and movement**

Each surviving cell's per-subject power change was correlated with the concurrent change in head-accelerometry movement vigor, and the spectra were parameterized into aperiodic (broadband 1/f) and periodic (oscillatory-peak) components. The whole-head high-beta power increase scaled with the movement increase ($\rho = 0.86$, robust to leave-one-out), but this coupling was carried by the broadband component alone: the oscillatory high-beta and gamma peaks showed no association with movement ($\rho = -0.06$ and $-0.22$, ns), whereas the aperiodic component increased during performance and showed a parallel, trend-level association with vigor at the scalp ($\rho = 0.73$) and source level (offset $\Delta = +0.15$, $p = 0.084$), reaching significance within several dipole clusters. The fast-band effects thus comprise a movement-robust oscillatory component and a broadband component that co-occurs with movement and cannot, at this sample size, be distinguished between movement-driven cortical activation and residual myogenic activity. The exploratory alpha suppression, by contrast, was movement-robust (a genuine oscillatory-peak reduction; $\rho = 0.08$). Because spectral parameterization at this sample size detects oscillatory peaks in only a minority of channel spectra, these decompositions are reported as sensitivity analyses rather than primary findings.

**Intra-brain: robustness and spatial distribution**

Two limits qualify the omnibus. Under leave-one-subject-out resampling the omnibus p-value ranged from 0.024 to 0.094, remaining below 0.05 in 4 of 12 folds (Supplementary Figure S4); the effect is present in the full sample but sensitive to individual participants. Second, the omnibus collapses spatial information: at the descriptive cell level, all 56 high-frequency network-by-band cells showed a positive change, with the largest effects in dorsal-attention–limbic coupling (high-beta $d_z = 0.85$) and visual-network couplings ($d_z \approx 0.73$–$0.75$) (Figure 3c). No individual network pair survived family-wise correction (all $p_{FWER} > 0.44$), so these locations are hypothesis-generating descriptions of where the distributed increase was strongest. Mirroring showed a similar but weaker pattern (54 of 56 cells positive); breathing did not (25 of 56).

**Intra-brain: age independence**

Adding age as a covariate left the performance effect essentially unchanged (adjusted $d_z = 0.61$, $p = 0.06$; age slope $p = 0.76$; connectivity–age $\rho = -0.01$). The slight increase in p relative to the unadjusted test ($p = 0.049$) reflects the degree of freedom spent on the covariate. Age was likewise unrelated to the (non-significant) mirror and breathing indices (both age slopes $p > 0.66$).

**Intra-brain: movement independence**

Each measure's performance-versus-baseline change was tested for association with concurrent head-accelerometry vigor, both as a rank correlation and after linear adjustment. The two leakage-robust measures were independent of movement: MIM change was uncorrelated with vigor change ($\rho = 0.43$, $p = 0.167$; adjusted vigor slope $p = 0.376$), and orthogonalized AEC — the largest effect ($dz = 1.85$) — showed no movement association ($\rho = 0.04$, $p = 0.897$; slope $p = 0.412$) (Supplementary Table S5). In contrast, coherence computed without leakage correction (aCOH) did track vigor ($\rho = 0.62$, $p = 0.031$); because aCOH is already flagged as susceptible to zero-lag leakage, this serves as a positive control. Convergently, AEC was comparably elevated during mirroring ($dz = 1.47$), a condition in which vigor did not increase relative to baseline, and was uncorrelated with vigor within that condition ($\rho = 0.32$, $p = 0.319$).

**Inter-brain: robustness and spatial distribution**

The Charlie Brown effect was robust to individual dyads, with both amplitude measures exceeding surrogate in every leave-one-dyad-out fold (envelope $d = 1.13$–$1.76$; orthogonalized $d = 0.97$–$1.81$). Spatially, the effect was broad and diffuse rather than localized: real coupling exceeded surrogate at more than 90% of the 1,024 cross-brain channel pairs during *Charlie Brown,* with no preference for homologous over heterologous electrode pairings and no concentration in a small subset of pairs (Figure 4, Panels E–F; per-condition and per-pair detail in Supplementary Figure S5 and Supplementary Table S8). Because the per-pair effects are descriptive and the pseudo-dyad surrogate is not fully independent, individual channel pairs are not interpreted. The estimates rest on few dyads ($n = 7$–$10$ per condition) and the surrogate reuses participants across pairings, so permutation p-values are approximate and effect sizes carry the primary interpretive weight. The composition of the ten dyads and their pseudo-dyad pairings is detailed in Supplementary Table S9.

**Inter-brain: age independence**

The high-frequency coupling was independent of age. Neither the high-beta nor the gamma increase was related to dyad mean age for either amplitude measure (Spearman $\rho = 0.31–0.35$, all $p > 0.15$), and both were essentially unchanged after age adjustment (Table 5, Panel C). Alpha-band coupling was the exception: it was present already at rest, scaled steeply with age ($\rho = 0.79$, $p = 0.01$), and did not survive age adjustment, confining the age relation to resting alpha rather than the high-frequency performance effect.
